# Optimization of somatic embryogenesis in *Euterpe edulis* Martius using auxin analogs and atomic force microscopy

**DOI:** 10.1101/2023.03.04.531114

**Authors:** Tamyris de Mello, Yanara dos Santos Taliuli, Tatiane Dulcineia Silva, Tadeu Ériton Caliman Zanardo, Clovis Eduardo Nunes Hegedus, Breno Benvindo dos Anjos, Edilson Romais Schmildt, Adésio Ferreira, Maicon Pierre Lourenço, Patricia Fontes Pinheiro, Glória Maria de Farias Viégas Aquije, José Carlos Lopes, Wagner Campos Otoni, Rodrigo Sobreira Alexandre

## Abstract

*Euterpe edulis* Martius is an endangered species of the Atlantic Forest, whose fruits have high antioxidant potential, and propagated exclusively by seeds. The present study assessed the ability of different auxin inducers and picloram analogs to trigger somatic embryogenesis in *E. edulis*. Immature seeds were harvested, and their zygotic embryos were excised and grown in MS culture medium supplemented with 2,4-D dichlorophenoxyacetic acid (2,4-D) or picloram at 150, 300, 450, 600 µM. The activity of picloram analogs triclopyr and clopyralid was evaluated in semisolid MS medium. At maturation and germination, picloram-derived calli and somatic embryos isolated from triclopyr-grown cultures were first transferred to pre-maturation medium and, after 30 days, to basal MS or MS medium supplemented with either 5 µM abscisic acid or 0.53 µM 1-naphthaleneacetic acid plus 12.3 µM 2-isopentenyladenine. Finally, somatic embryos with root protrusions were transferred to MS medium devoid of sucrose for 30 days and then acclimatized ex vitro. Scanning, transmission, and atomic force microscopy revealed that picloram was superior to 2,4-D but less effective than triclopyr (100 µM) in inducing embryogenesis. Maturation and germination of somatic embryos in E. edulis can be maximized by 5 µM abscisic acid, and selecting calli via atomic force microscopy.

**Highlight:** This work opens novel roads for embryogenic induction, using a new and more efficient inducer than the usual ones, and an innovative evaluation technique based on AFM.

## INTRODUCTION

*Euterpe edulis* Martius is a species native to the Atlantic Forest. Its fruits are rich in bioactive compounds and nutrients, including antioxidants such as anthocyanins and phenolic acids. However, the conservation of this species is hindered by deforestation and illegal palm heart extraction (Schulz *et al*., 2016).

The propagation of *E. edulis* occurs exclusively through seeds, and tillering is not possible because no shoots are produced at the base of the plant (Schulz *et al*., 2016). Crucially, owing to the strong recalcitrance of the seeds, germination is slow and uneven, and can take up to 60–90 days (Cursi and Cicero, 2014).

Somatic embryogenesis is a promising in vitro propagation technique for palm trees (Campos *et al*., 2020). With the oil palm *Elaeis guineenses* Jacq., it has resulted in uniform material for planting programs and genetic improvement (Beulé *et al*., 2011). Embryogenic calli induced from palm tree explants can be used as target tissues in genome-editing protocols (Yarra *et al*., 2020).

The induction and development of somatic embryos depends on culture medium composition, explants, genotypes, inducers, and their concentrations (Rose *et al*., 2010). Zygotic embryos are the most commonly used explants for somatic embryogenesis in palm trees, particularly those of the genus *Euterpe* (Ferreira *et al*., 2022a; Freitas *et al*., 2016; Guerra and Handro, 1998; Oliveira *et al*., 2022; Scherwinski-Pereira *et al*., 2012). They are highly responsive during in vitro culture and represent a valuable tool in somatic embryogenesis of economically important species, research on adult tissue, and cloning of genetically improved plants (Silva-Cardoso *et al*., 2020).

Picloram (4-amino-3,5,6-trichloropyridine-2-carboxylic acid) is a synthetic pyridinecarboxylic acid herbicide (Marco-Brown *et al*., 2014) and an auxin mimic. Owing to its efficient absorption and mobilization, as well as its rapid metabolism, picloram has been used succesfully in somatic embryogenesis (Karun *et al*., 2004). While picloram has yielded good results in palm trees, such as *Bactris gasipaes* (Steinmacher *et al*., 2007; Valverde *et al*., 1987), *Calamus merrillii* Becc., *Calamus subinermis* Wendl. ex Becc. (Goh *et al*., 1999; 2001), *Euterpe oleracea* (Freitas *et al*., 2016; 2018; Scherwinski-Pereira *et al*., 2012), and *E*. *edulis* (Mello *et al*., 2023; Oliveira *et al*., 2022), its analogs, such as fluroxypyr (4-amino-3,5-dichloro-6-fluoro-2-pyridyloxyacetic acid), triclopyr (3.5,6-trichloro- 2-pyridinyloxyacetic acid), clopyralid (3,6-dichloro-pyridine-2-carboxylic acid), and aminopyralid (4- amino-3,6-dichloropyridine-2-carboxylic acid), have not been tried for this purpose.

Traditionally, the embryogenic development of calli has been documented using optical, scanning, and transmission electron microscopy (Ferreira *et al*., 2022a; Freitas *et al*., 2016; Oliveira *et al*., 2022; Scherwinski-Pereira *et al*. 2012; Silva-Cardoso *et al*., 2020).

Developed for the first time in 1986 (Binnig *et al*., 1986), atomic force microscopy (AFM) differs from other imaging techniques (Demir-Yilmaz *et al*., 2021), as it is based on attractive and repulsive forces between a probe and a sample. Essentially, the probe deflection is a function of the interaction with the sample. AFM has become an important tool in material science (Dias *et al*., 2020; Last *et al*., 2010; Pillet *et al*., 2014; Xiao and Dufrêne, 2016) as it benefits from a nanometer resolution, low invasiveness, simplicity, and rapidity of manipulation (Hu *et al*., 2022).

Given the good results obtained with picloram, we hypothesized that its analogs triclopyr and clopyralid could be promising compounds for somatic embryogenesis in *E. edulis*, and that AFM could help in the identification of embryogenic calli. Hence, in the present study, we assessed somatic embryogenesis of *E. edulis* treated with different auxin inducers and picloram analogs.

## MATERIALS AND METHODS

### Collection and disinfection

The fruits of *E. edulis* were harvested from a plant with a height of 6.7 m and diameter at breast height (-1.30 m) of 13.4 cm. The tree was located near the Parque Nacional do Caparaó, ES, Brazil, at 20°32’45.4’’ S, 41°49’35.7’’ W, and an altitude of 951 m. The fruits were selected for their high level of anthocyanin in the pulp, harvested approximately 168 days after anthesis, and sent to the Laboratory of Forest Seeds and Plant Tissue Culture at the Center for Agricultural Sciences and Engineering, Federal University of Espírito Santo (UFES).

After cleaning the fruits with water and neutral detergent, the tegument was removed, and the fruits were immersed in a 2% ascorbic acid solution to prevent phenolic oxidation. Using a laminar flow chamber, *E. edulis* seeds were immersed in 70% ethyl alcohol for 1 min, 2.5% sodium hypochlorite (Candura) for 10 min, and amoxicillin solution (Germed) at 3 g L^-1^ for 10 min. After each step, the seeds were washed three times in distilled and autoclaved water.

### Induction

Zygotic embryos were extracted from immature seeds and used in experiments I and II. In experiment I, the embryos were arranged in glass Petri dishes (90 × 15 mm) and immersed in 6 mL stationary MS liquid culture medium (Murashige and Skoog, 1962) (Sigma) containing 0.1 g L^-1^ myo- inositol (Sigma), 30 g L^-1^ sucrose (Dinâmica), and 1 g L^-1^ polyvinylpyrrolidone (PVP) (Synth) at pH 5.7 ± 0.1. The cultures were supplemented with 2,4-dichlorophenoxyacetic acid (2,4-D) (Sigma) and picloram (Sigma) at 150, 300, 450, and 600 µM. In experiment II, 5.5 g L^-1^ agar-agar (Dinâmica) was added and the culture was supplemented with picloram or its analogs triclopyr (Sigma) and clopyralid (Sigma) at 100, 125, 150, and 175 µM. All growth inducers were filter-sterilized and added to autoclaved culture medium.

The cultures were maintained in a growth room at 27 ± 2°C in the dark for 170 days. In experiment I, oxidation (%), calli (%), embryogenic calli (%), total callus area (mm²), embryogenic area (mm²), and embryogenic percentage were analyzed. In experiment II, oxidation (%), callogenesis (%), induction rate (%), callus mass (g), and total callus area (mm²) were measured.

#### Scanning electron microscopy

Calli and embryos obtained after induction were fixed in Karnovsky solution, dehydrated in an ethylic series to absolute alcohol, dried to a critical point with CO_2_ (Autosamdri 815; Tousimis), arranged in stubs, and subjected to gold sputtering (Desk V; Denton Vacuum).

Samples were analyzed with a scanning electron microscope (JSM-6610LV; Jeol) at the Laboratory of Cellular Ultrastructure Carlos Alberto Redins (LUCCAR), UFES.

#### Histological and structural analyses

For anatomical studies, samples were dehydrated in an ethanol series and soaked in methacrylate (Historesin^®^, Leica). Transverse and longitudinal sections (5 μm in thickness) were obtained with an automatic rotating microtome (RM2155; Leica) equipped with a disposable glass knife, and stained with toluidine blue at pH 4.0 (O’Brien and McCully, 1981) for 15 min. The slides were mounted in synthetic resin (Permount^®^; Fisher Scientific) and analyzed under a light microscope (Olympus-AX 70) coupled with a photomicrography system (Olympus U-Photo) at the Laboratory of Plant Anatomy, Department of Plant Biology, Federal University of Viçosa.

#### Transmission electron microscopy

Samples from different experimental groups were collected and fixed in Karnovsky’s solution containing 2.5% glutaraldehyde, 2% paraformaldehyde, and 0.1 M cacodylate buffer. Subsequently, the tissues were post-fixed in osmium tetroxide, dehydrated in ethanol (30%, 50%, 70%, 90%, and 100%), and embedded in epoxy resin (Embed 812; Electron Microscopy Sciences). Ultrathin sections (60–80 nm in thickness) were obtained using an ultramicrotome (UCT; Leica Microsystems) before staining with uranyl acetate and lead citrate. Representative electron micrographs of different groups were obtained using a transmission electron microscope (JEM-1400; JEOL), operated at 120 kV and 12000× magnification, at LUCCAR-UFES.

#### Atomic force microscopy

Samples of nodular calli measuring approximately 0.5 cm^2^ were dehydrated in an ethylic series to absolute alcohol, dried to a critical point with CO2, and placed on glass slides using enamel as glue.

AFM images were collected using an alpha300 R confocal microscope (WITec GmbH) operated in non-contact mode, with a Si_3_N_4_ cantilever, a nominal constant of 42 N m^-1^, resonant frequency of ∼285 kHz, scan rates of 0.3–1.0 Hz, and scan range of 2500–10000 nm.

Phase and light microscopy images were collected along with topographical images. Phase images were used to estimate the physical properties of the calli, including hardness, adhesion, and viscoelasticity.

Surface asymmetry (SSK) was obtained from Equation 1 and the peak-to-peak height, given by the difference between the highest and lowest peak heights, was used to assess surface roughness.

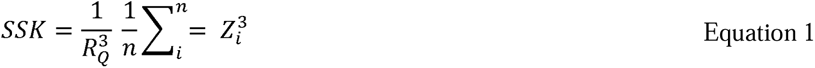

Zi is the height at position I, R_Q_ is the mean square of the height deviation, and n is the number of points on the image grid.

In general, SSK = 0 suggests a symmetrical or even data distribution around the median plane; whereas SSK ≠ 0 suggests an asymmetrical distribution, consisting of either a flat surface with small peaks (SSK > 0) or small valleys (SSK < 0).

Kurtosis (SKU) was determined by Equation 2 using WITec software. It indicates whether the data are arranged horizontally or perpendicularly to the mean.

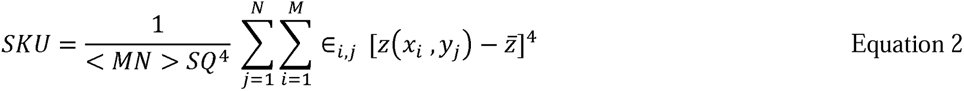

M and N are the numbers of data points in X and Y, respectively, SQ is the root mean square deviation, and Z is the height of the surface relative to the mean plane. SKU > 3.00 indicates the presence of excessively high peaks or deep valleys, SKU < 3.00 indicates surfaces with protruberances, and SKU = 3 corresponds to a normally distributed surface height.

Analyses were performed at the Laboratory for Research and Development of Methodologies for Oil Analysis, Department of Chemistry, UFES.

#### Chemical analysis of inducers

Statistical analyses were performed using Python 3.x. Descriptors of the four molecules (2,4-D, picloram, triclopyr, and clopyralid) were obtained from the RDKIT library (http://www.rdkit.org/), which uses the SMILES method (https://pubs.acs.org/doi/abs/10.1021/ci00057a005) for molecular representation. Principal components analysis (PCA) was carried out using the scikit-learn library (https://dl.acm.org/doi/10.5555/1953048.2078195). Graphs were generated using Matplotlib (Hunter, 2007), along with Numpy (Harris, 2020).

#### Experimental design

The experiments followed a completely randomized design, based on either a 2 × 4 factorial scheme (inducers I: 2,4-D and picloram × concentrations: 150, 300, 450, and 600 µM) or a 3 × 4 factorial scheme (inducers II: picloram, triclopyr, and clopyralid × concentrations: 100, 125, 150, and 175 µM). Each experiment consisted of four replicates with five zygotic embryos. Data were subjected to analysis of variance, F test between inducers in Experiment I, mean test between inducers in Experiment II, Tukey’s test (*P < 0.05*), and regression analysis using R software (R Core Team, 2020).

### Maturation and germination

All embryogenic calli induced in Experiment II were transferred to pre-maturation MS medium supplemented with 0.1 g L^-1^ myo-inositol and 30 g L^-1^ sucrose, without the addition of regulators. The pH was adjusted to 5.7 ± 0.1 before adding 5.5 g L^-1^ agar-agar. The cultures were kept in a growth room at 27 ± 2°C, in the dark, for 30 days, and the number of somatic embryos was counted.

Next, embryogenic calli induced with picloram (125, 150, and 175 µM) were transferred to MS medium containing 0.1 g L^-1^ myo-inositol and 30 g L^-1^ sucrose. The pH was adjusted to 5.7 ± 0.1 before adding 5.5 g L^-1^ agar-agar. After autoclaving, the medium was supplemented with 5 µM abscisic acid (ABA) or 0.53 µM 1-naphthaleneacetic acid (NAA) and 12.3 µM 2-isopentenyladenine (2-iP) (Guerra and Handro, 1998). Embryogenic calli obtained upon induction with triclopyr (100 µM) were arranged in the same culture medium as above, kept in a growth room at 27 ± 2°C in the dark until root protrusion, and the percentage of germination was evaluated.

### Growth and acclimatization of somatic seedlings

Somatic embryos displaying root protrusions and previously induced with triclopyr (100 µM) were transferred to MS medium containing 0.1 g L^-1^ myo-inositol, 30 g L^-1^ sucrose, and 1 g L^-1^ PVP. The pH was adjusted to 5.7 ± 0.1 before adding 4 g L^-1^ agar-agar. The cultures were kept in a growth room at 27 ± 2°C, with a photoperiod of 8/16 h and irradiance of 30 µmol m^-2^ s^-1^ until obtaining emblings of approximately 5 cm, with two expanded leaves and at least two roots.

Emblings were transferred to MS culture medium containing 0.1 g L^-1^ myo-inositol, 1 g L^-1^ PVP, and 4 g L^-1^ agar-agar. After 30 days, the tubes with the emblings were transferred to room temperature, and 7 days later they were placed in an ex vitro environment consisting of plastic cups (590 mL) containing autoclaved substrate (TerraNutri^®^) moistened with distilled and autoclaved water. Each cup was covered with another cup (300 mL) to maintain the humidity during the first 7 days. Following that period, two holes (Ө = 0.5 cm) were made in the upper cup every 2 days, and the cultures were incubated for another 10 days.

## RESULTS

### Induction

#### Experiment I

*E. edulis* zygotic embryos (Fig.. 1a–d) treated with 2,4-D appeared strongly oxidized even before callus formation (Fig.. 1e); this was not the case with picloram-induced embryos (Fig.. 1f–i). The low callus percentage of 2,4-D-treated embryos (Fig.. 2a, b) was matched by 100% oxidation at 450 and 600 µM 2,4-D (Fig.. 2c). Instead, zygotic embryos treated with picloram achieved 100% callus formation (Fig.. 2a, b), formed embryogenic regions and somatic proembryos, and presented much lower oxidation (Fig. 2a–c).

**Fig. 1.**
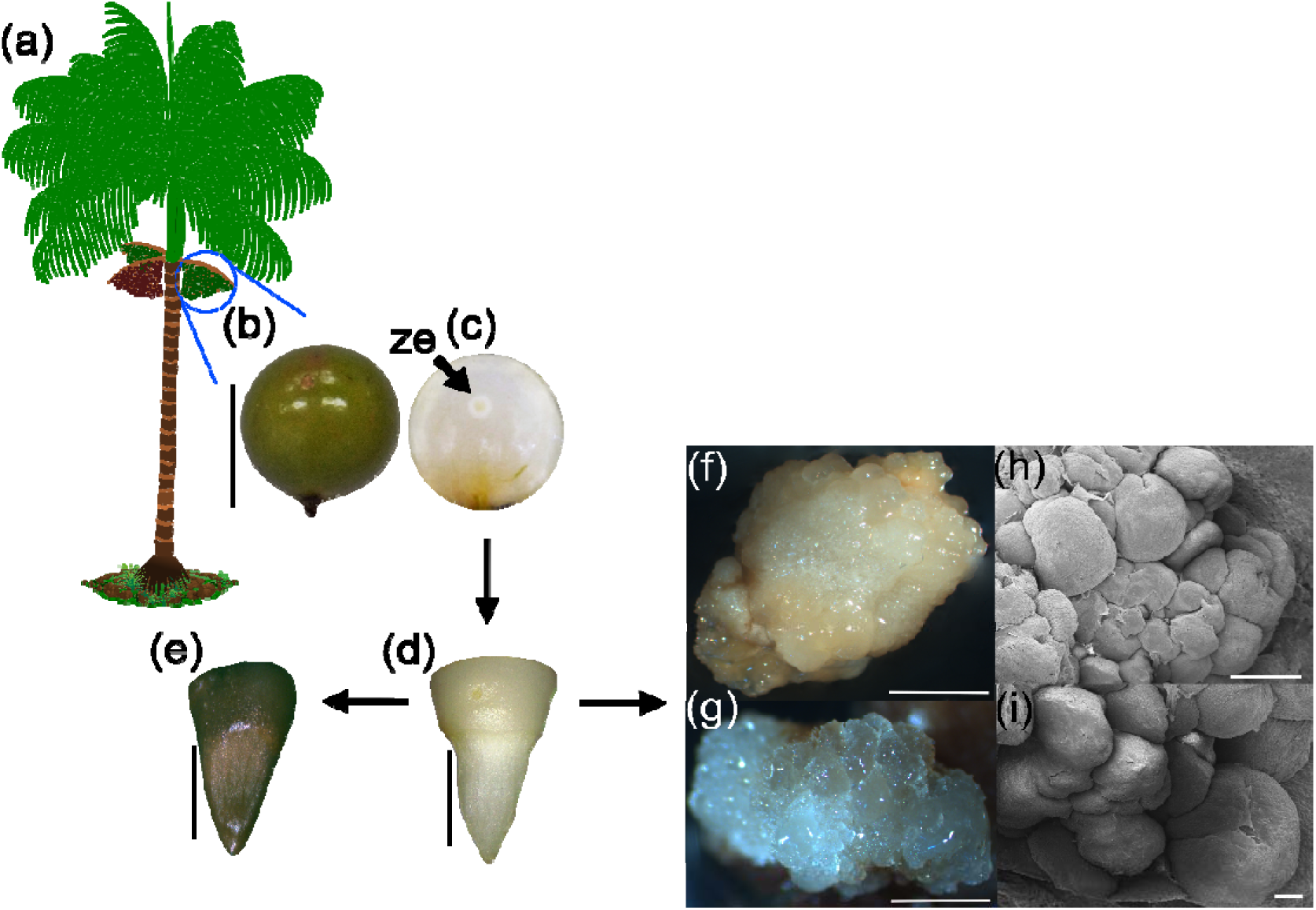
Embryogenesis in *Euterpe edulis*. (a) Schematic llustration of a fruiting *E. edulis* tree. (b–i) ≈180 days after anthesis (ze. zygotic embryo) (c), zygotic embryo detached from the seed (d), oxidized zygotic embryo (e), embryogenic callus treated with 150 µM picloram (f), embryogenic area (g), and a magnification of the embryogenic area (h, i). Bar: 1.0 cm (a); 5000 µm (f), 2000 µm (d, e, g), 1000 µm (h), and 200 µm (i).

**Fig. 2.**
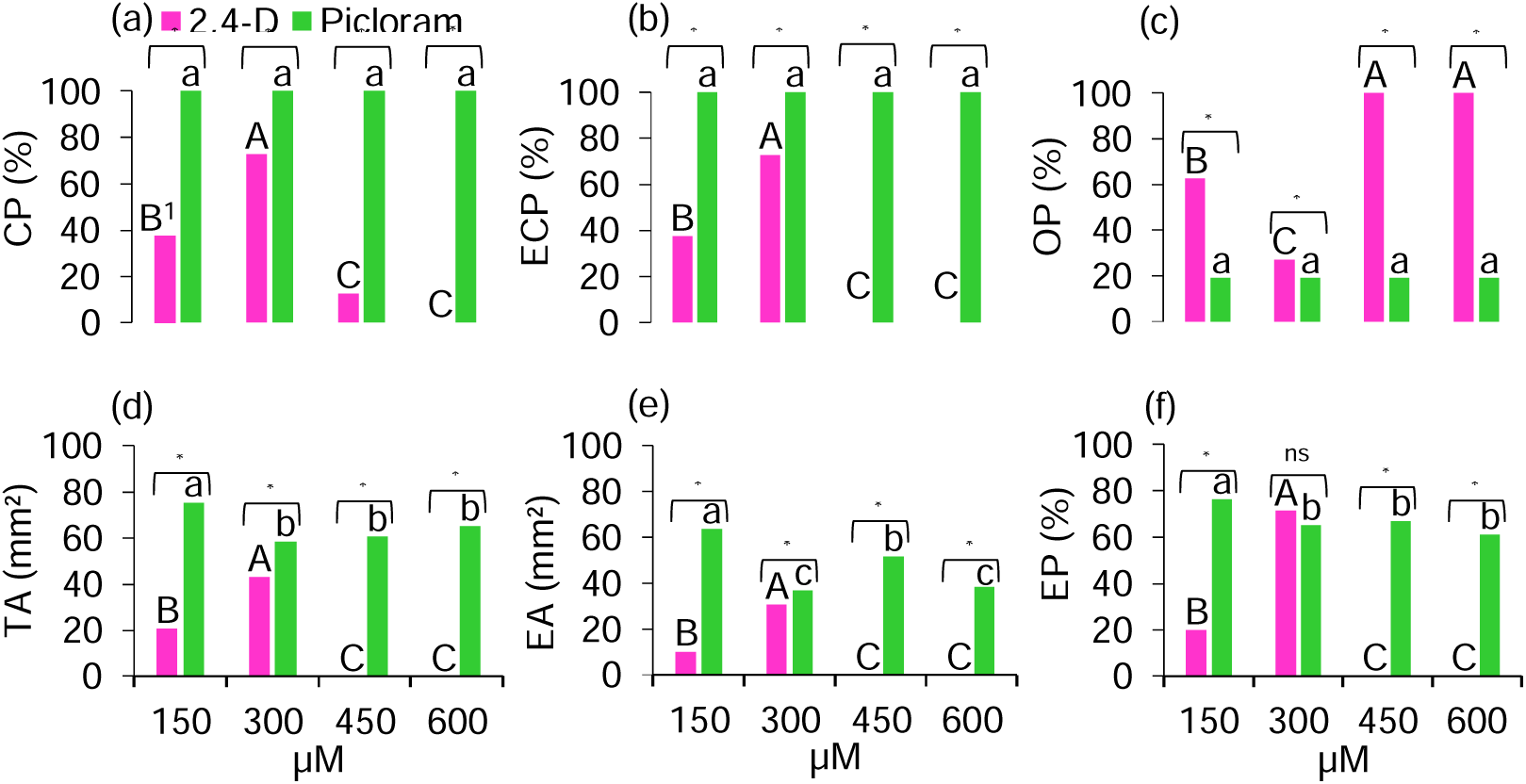
Induction of somatic embryogenesis in immature zygotic embryos of *E. edulis* upon treatment with 2,4-D or picloram. (a) Callus percentage (CP), (b) embryogenic callus percentage (ECP), (c) oxidation percentage (OP), (d) callus total area (TA), (e) embryogenic area (EA), and (f) percentage of embryogenic area (EP). ^1^Averages followed by the same capital letter (2,4-D, pink bar) and lowercase letter (picloram, green bar) between concentrations do not differ from each other according to Tukey’s test (*P < 0.05*). Differences between the two inducers were deemed *significant (*P < 0.05*) or ns, not significant (F test).

No adjustment for these variables was necessary in the regression model. A significant interaction between all variables was detected with respect to both embryogenic inducer and concentration (Fig. 2). Picloram led to the highest mean callus percentage, embryogenic callus percentage, total callus area, callus embryogenic area, and embryogenic percentage, but the lowest oxidation percentage.

#### Experiment II

Zygotic embryos treated with picloram and triclopyr produced both non-embryogenic and embryogenic calli (Fig. 3a–e). With both inducers, the calli presented embryogenic regions (Fig. 3c, e) and, in the case of triclopyr, somatic embryos after 170 days (Fig. 3e). Instead, most zygotic embryos treated with clopyralid gave rise to non-embryogenic structures and abnormal seedlings (Fig. 3f, g).

**Fig. 3.**
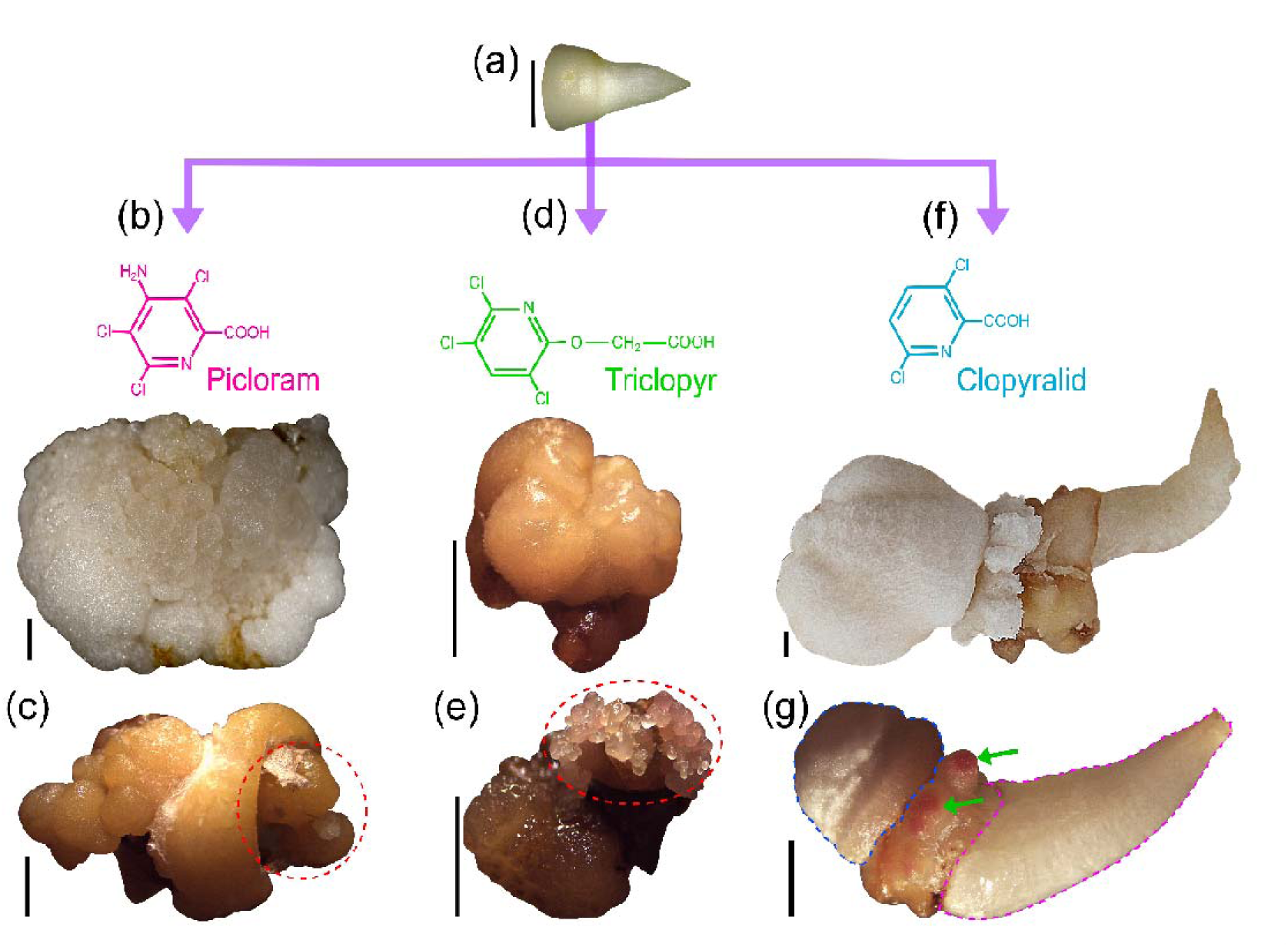
Appearance of calli obtained after induction of a zygotic embryo of *Euterpe edulis* with picloram, triclopyr or clopyralid. (a) Detail of an isolated zygotic embryo used as an explant for embryogenic induction. (b, c) Non-embryogenic (b) and embryogenic (c) calli obtained following induction with picloram; the dashed red line highlights the embryogenic area. (d, e) Non-embryogenic (d) and embryogenic (e) calli obtained following induction with triclopyr; the dashed red line highlights the embryogenic area with somatic embryos. (f, g) Non-embryogenic structures (f), abnormal seedling with haustorium (dashed blue line), roots (green arrows), and caulicle (dashed pink line) (g) obtained following induction with clopyralid. Bar: 2000 µm (a–g).

After 170 days, somatic proembryos appeared in picloram-, triclopyr-, and clopyralid-treated calli, albeit at different proportions (Fig.s 4–7). Picloram-induced calli were larger than those induced with triclopyr.

**Fig. 4.**
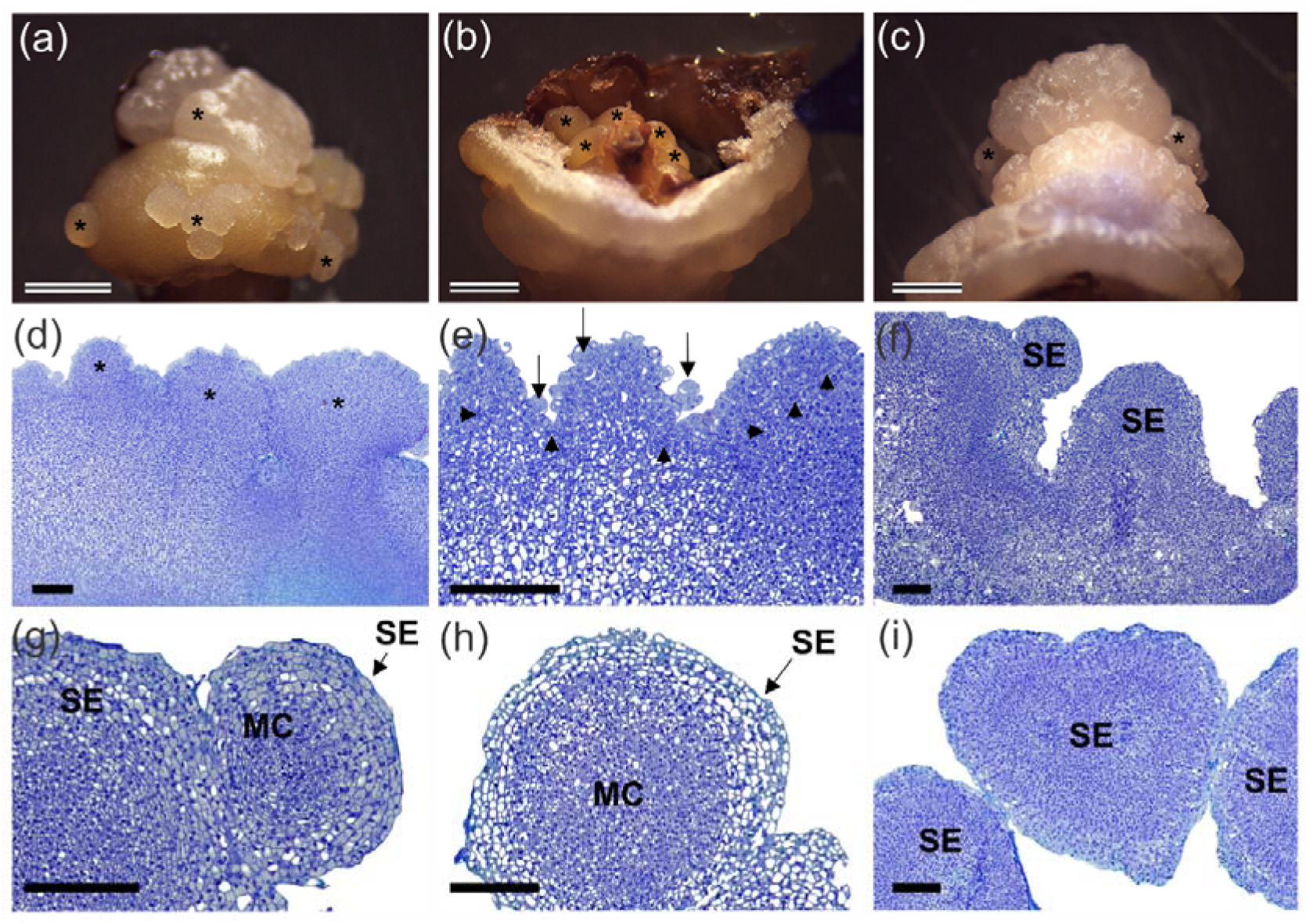
Somatic embryos obtained from embryogenic calli of *Euterpe edulis* induced with picloram. (a–c) Embryogenic calli, highlighting the formation of somatic embryos (*). (d–i) Histological sections of somatic proembryos (d), subepidermal cells forming clusters of anticlinal and periclinal dividing cells (arrows) (e), somatic embryos (SE) (f), somatic embryos and meristematic cells (MC) (g, h), and individual somatic embryo (i). Bar: 2000 µm (a–c) and 400 µm (d–i).

**Fig. 5.**
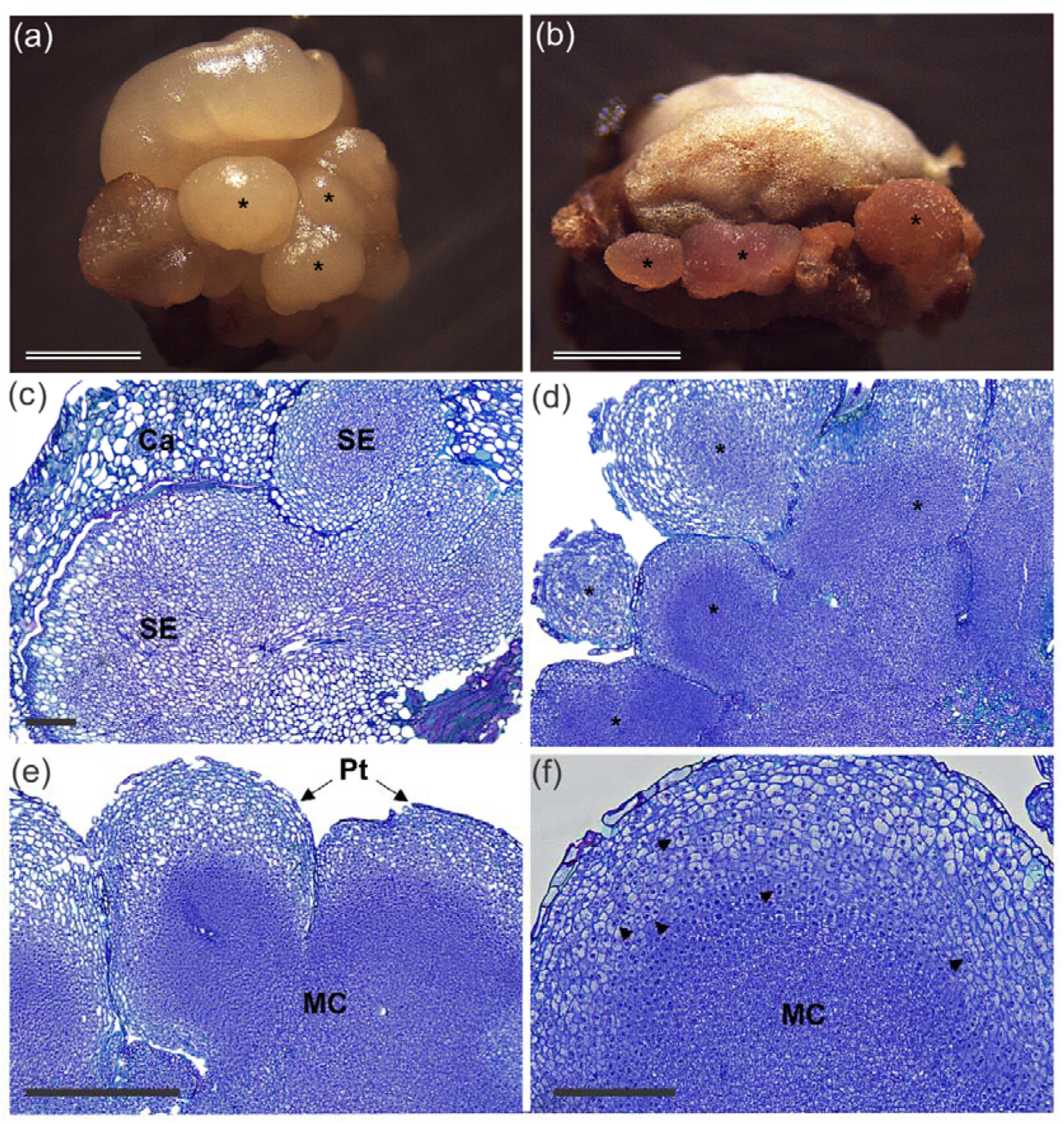
Somatic embryos obtained from embryogenic calli of *Euterpe edulis* induced with triclopyr. (a, b) Embryogenic calli, highlighting the regions in which somatic embryos are being formed (*). (c–f) Histological sections of somatic proembryos (SE) surrounded by callus (Ca) (c), somatic embryos (*) (d), meristematic cells (MC) and protoderm (Pt) (e), and subepidermal cells in anticlinal and periclinal division (arrows) (f). Bar: 2000 µm (a, b) and 400 µm (c–f).

**Fig. 6.**
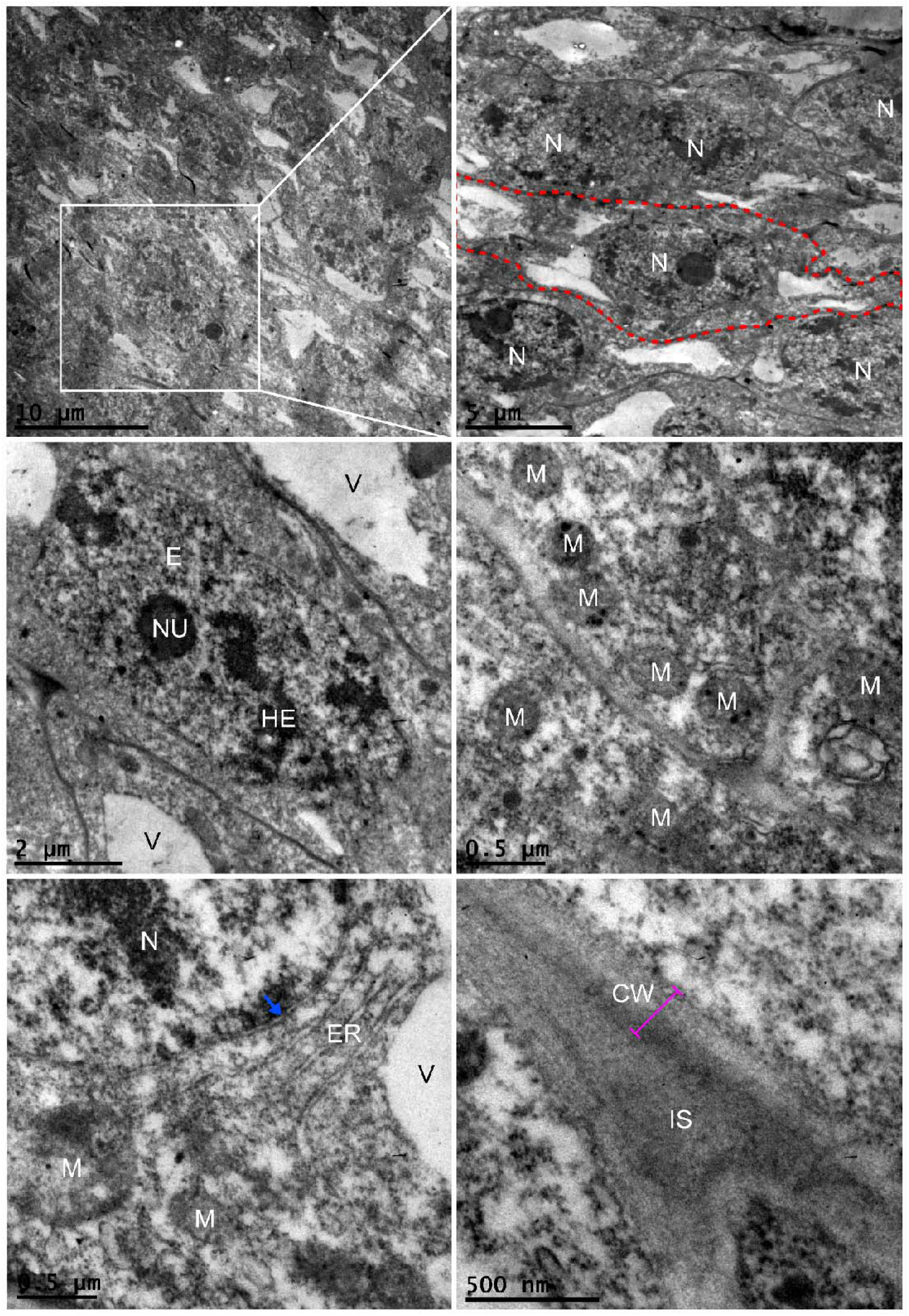
Ultrastructure of embryogenic calli induced with triclopyr (100 µM). (a) Embryogenic tissue sample at low magnification. (b–f) Detail of the embryogenic callus showing a cell (dashed red line) and nuclei (N) (b); nucleolus (NU), heterochromatin (HE), euchromatin (E), and vacuoles (V) (c); mitochondria (M) (d); nucleus and nuclear membrane (blue arrow), endoplasmic reticulum (ER), and mitochondria (e); and cell wall (CW) with intercellular space (IS) (f).

**Fig. 7.**
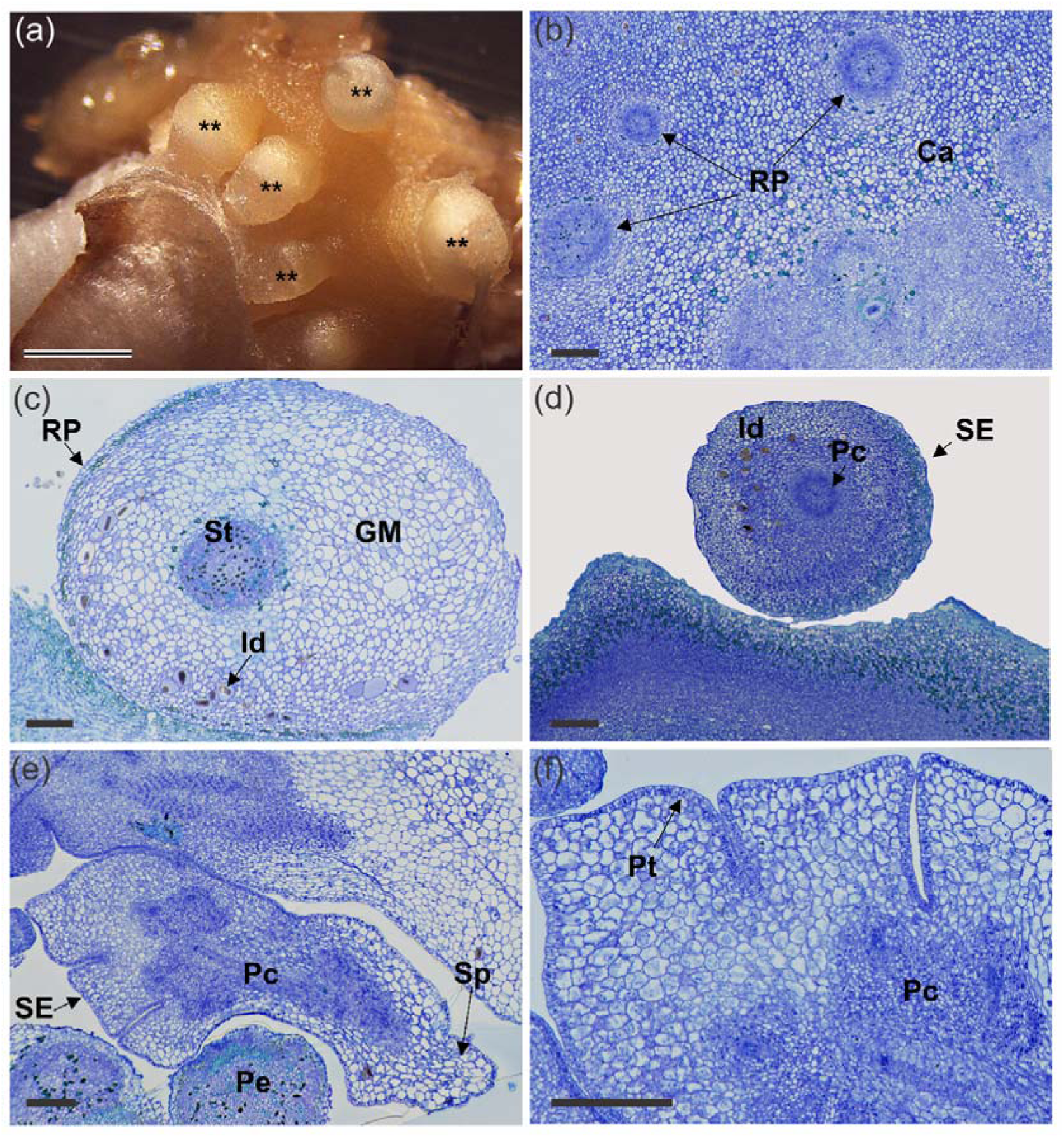
Somatic embryos obtained from embryogenic calli of *Euterpe edulis* induced with clopyralid. (a) Embryogenic calli, highlighting the regions in which root primordia are being formed (**). (b–f) Histological sections with root primordia (RP) surrounded by callus (Ca) (b); vascular tissue (St), ground meristem (GM), and idioblasts (Id) (c); globular somatic embryo (SE) with procambium (Pc) idioblasts (d); proembryo (Pe) and coleoptilar somatic embryo with procambium and primary root (Sp) (e); and somatic embryo with procambium and protoderm (Pt) (f). Bar: 2000 µm (a) and 400 µm (b–f).

After 150 days of induction, meristematic cells and proembryos were observed in embryogenic calli. Following histological sectioning, layers of intensely stained meristematic cells were observed. These regions differed from those of the embryogenic callus because they contained clusters of small isodiametric cells characterized by dense cytoplasm and conspicuous nuclei, as well as areas of intense cell division corresponding to meristematic centers.

Histological sections revealed subepidermal cells forming clusters in both anticlinal and periclinal division planes (Fig. 4e and Fig. 5f) and intensely dividing meristematic zones (Fig. 4f–i and Fig. 5e, f). Several cells in the central region of the embryo displayed an elongated shape, characteristic of cells of the procambium (Fig. 4f, i; Fig. 5e; and Fig. 6d–f). In addition to the procambium, somatic embryos presented a defined protoderm (Fig. 5e and Fig. 6f) and bipolarity in more advanced stages (Fig. 4i and Fig. 6e). Histological analysis revealed the absence of vascular attachment of the somatic embryo to the callus or a connection between the embryo’s base and the tissue of origin (Fig. 4g–i and Fig. 6d, e).

Embryogenic calli induced with triclopyr (100 µM) harbored cells with clear embryogenic characteristics, such as dense cytoplasm, evident nuclei and nucleoli, large and electron-dense, irregularly shaped nuclei, higher nucleus/cytoplasm ratio, fragmented vacuoles (Fig. 6a, b), lower heterochromatin-to-euchromatin content, numerous mitochondria, visible endoplasmic reticulum (Fig. 6c–e), cell components closely linked to the cell wall, and thick cell walls with visible intercellular space (Fig. 6f).

Clopyralid-induced calli presented instead small embryogenic areas with only a few somatic embryos. Most zygotic embryos formed abnormal seedlings, in which the growth of roots (Fig. 7a) was confirmed by the presence of root vascular bundles (Fig. 7b, c). Many idioblasts with raphides were also observed (Fig. 7c, d).

A significant interaction was observed between the inducers and their concentrations with respect to proembryo number, callus mass, and callus area. The induction rate showed a significant difference between inducers and concentrations; whereas callogenesis and oxidation displayed a significant difference only across inducers (Table 1).

**Table 1.**
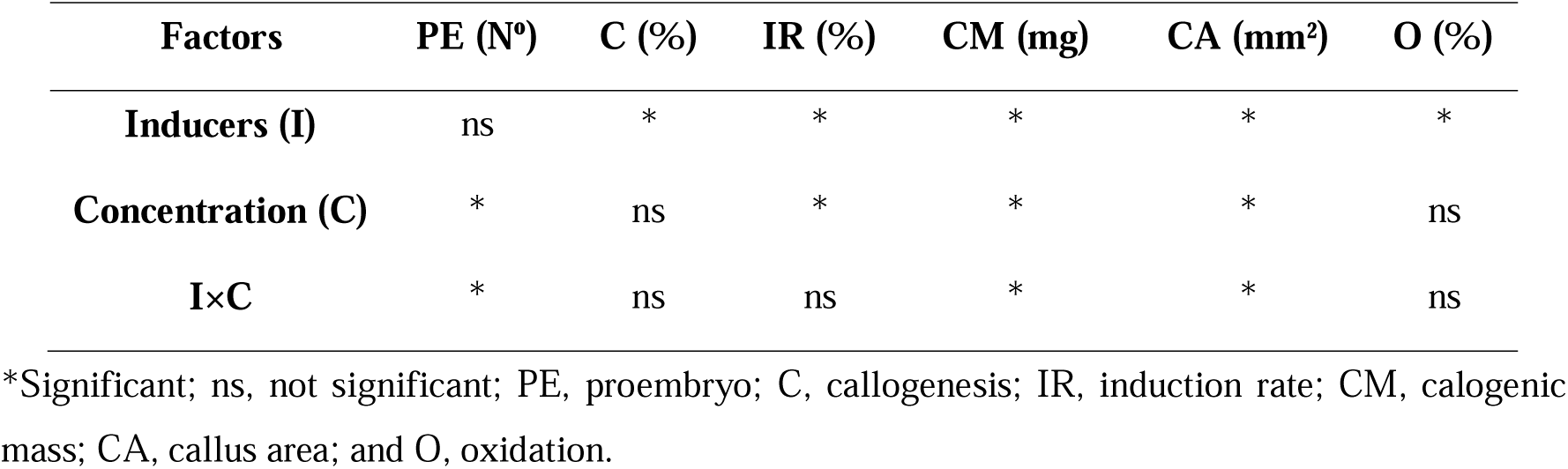
Significance of each variable within each factor (*P < 0.05*).

Triclopyr (100 µM) led to the highest number of proembryos (12) compared to other inducers (Fig. 8a), but yielded the lowest callus mass and callus area at all concentrations tested (Fig. 8b, c). Although clopyralid resulted in the highest mean callus mass and callus area, the structures were non-embryogenic and derived from abnormal seedlings.

**Fig. 8.**
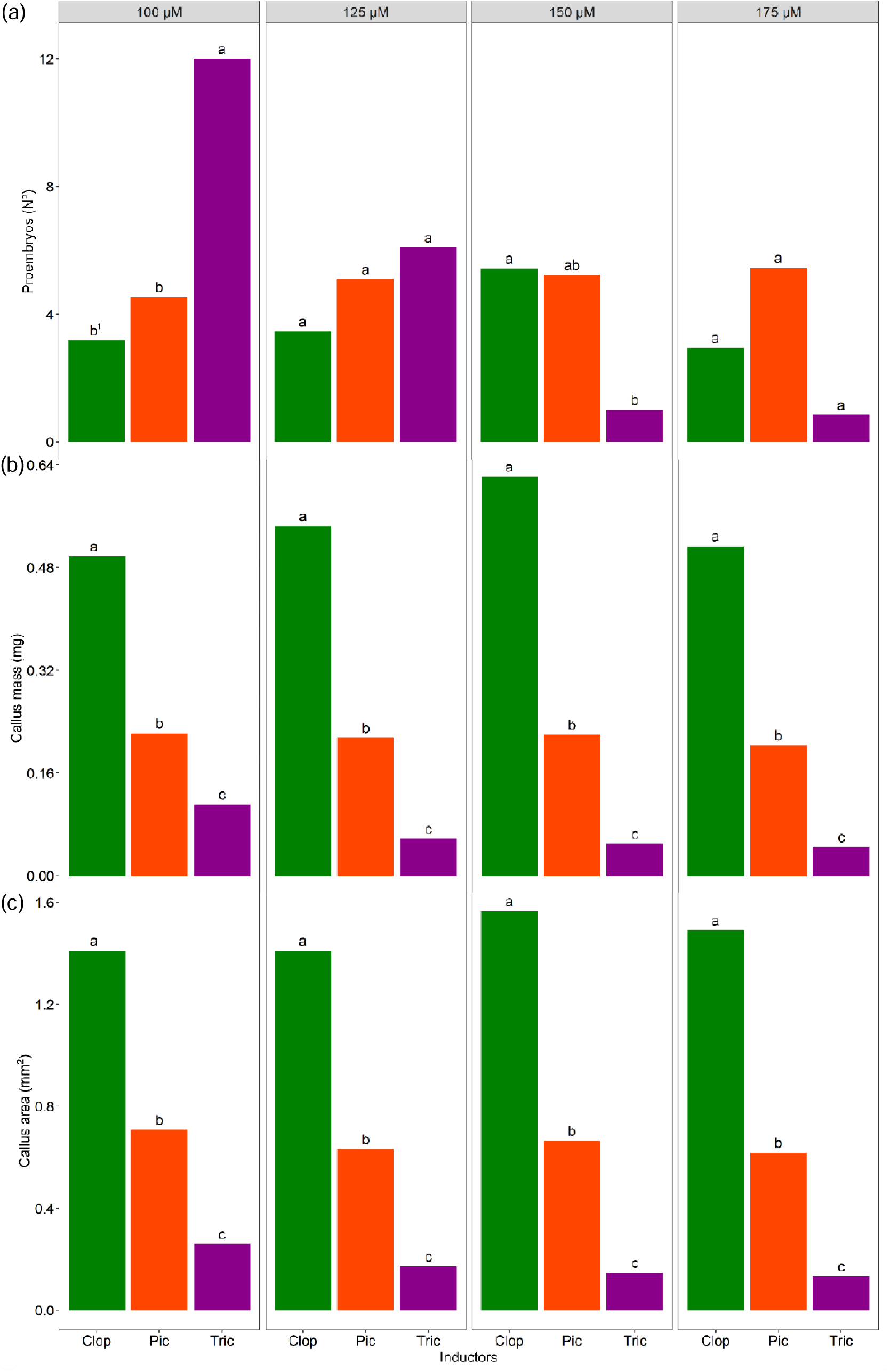
Embryogenic parameters in *Euterpe edulis* after 170 days of induction with clopyralid (Clop), picloram (Pic), and triclopyr (Tric) at different concentrations (100, 125, 150, and 175 µM). (a) Number of somatic embryos, (b) callus mass, and (c) callus area. ^1^Means not followed by the same letter in the comparison between inducers differ according to Tukey’s test (*P < 0.05*).

The three inducers yielded completely different results (Fig. 9). The number of proembryos and the callogenic mass did not differ across picloram concentrations, with a mean of 5.07 proembryos and 0.2137 g, respectively (Fig. 9a, b); while callus area showed only a slight decrease with increasing picloram concentration (Fig. 9c). In contrast, triclopyr led to a dramatic drop in the number of proembryos, as well as a gradual decline in callogenic mass and area with increasing concentrations (Fig. 9a–c). Finally, clopyralid had no effect on the number of proembryos (Fig. 9a), but displayed a quadratic behavior with respect to callogenic mass and area at increasing clopyralid concentrations (Fig. 9b, c).

**Fig. 9.**
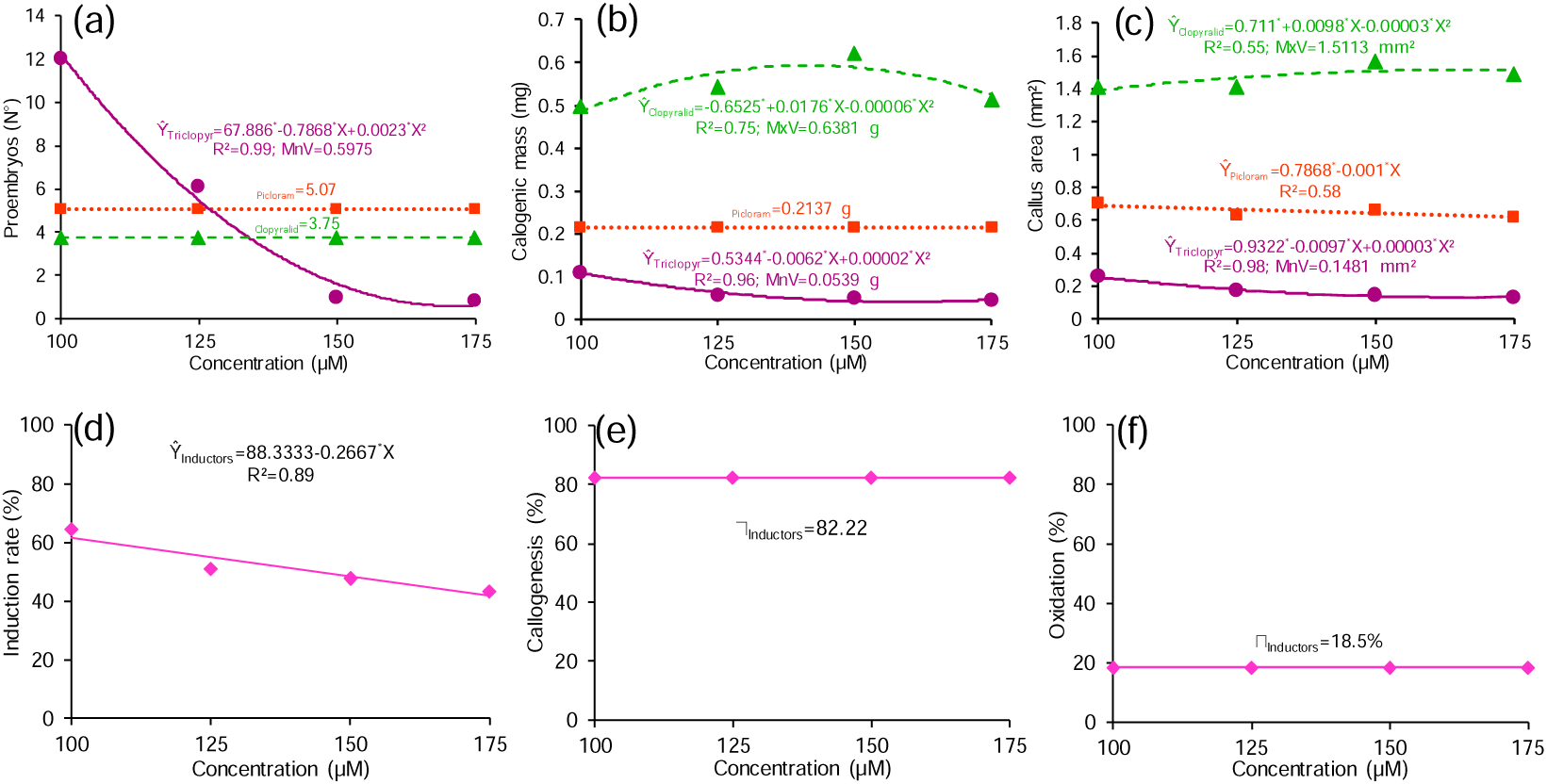
Embryogenic parameters of *Euterpe edulis* after 170 days of induction with clopyralid (green line), triclopyr (purple line), and picloram (orange line) at different concentrations (100, 125, 150, and 175 µM). (a) Number of somatic embryos, (b) callus mass, (c) callus area, (d) induction rate, (e) callogenesis, and (f) oxidation. *Statistically significant coefficient at 5% by the *t*-test. MxV, maximum value; MnV, minimum value.

The induction rate decreased linearly with increasing concentrations of inducers; whereas callogenesis and oxidation were not altered by the concentration of inducer, averaging 82.22% and 18.5%, respectively (Fig. 9d–f).

Clopyralid and picloram achieved the highest callogenesis and induction rates (Fig. 10a, b). In contrast, triclopyr displayed the lowest callogenesis and induction rates, which could be explained by the highest degree of explant oxidation (Fig. 10c).

**Fig. 10.** Callogenesis, induction rate, and oxidation of *Euterpe edulis* after 170 days of induction with clopyralid (Clop), picloram (Pic), and triclopyr (Tric) at different concentrations (100, 125, 150, and 175 µM). (a) Callogenesis, (b) induction rate, and (c) oxidation of explant. ^1^Means not followed by the same letter in the comparison between the inducers differ according to Tukey’s test (*P < 0.05*).

Picloram induced the formation of differently shaped and colored calli. These included non-embryogenic opaque white rough calli, spongy white rough calli, white smooth calli, and yellow smooth calli (Fig. 11a–h); as well as embryogenic rough yellow calli (Fig. 11i, j).

**Fig. 11.**
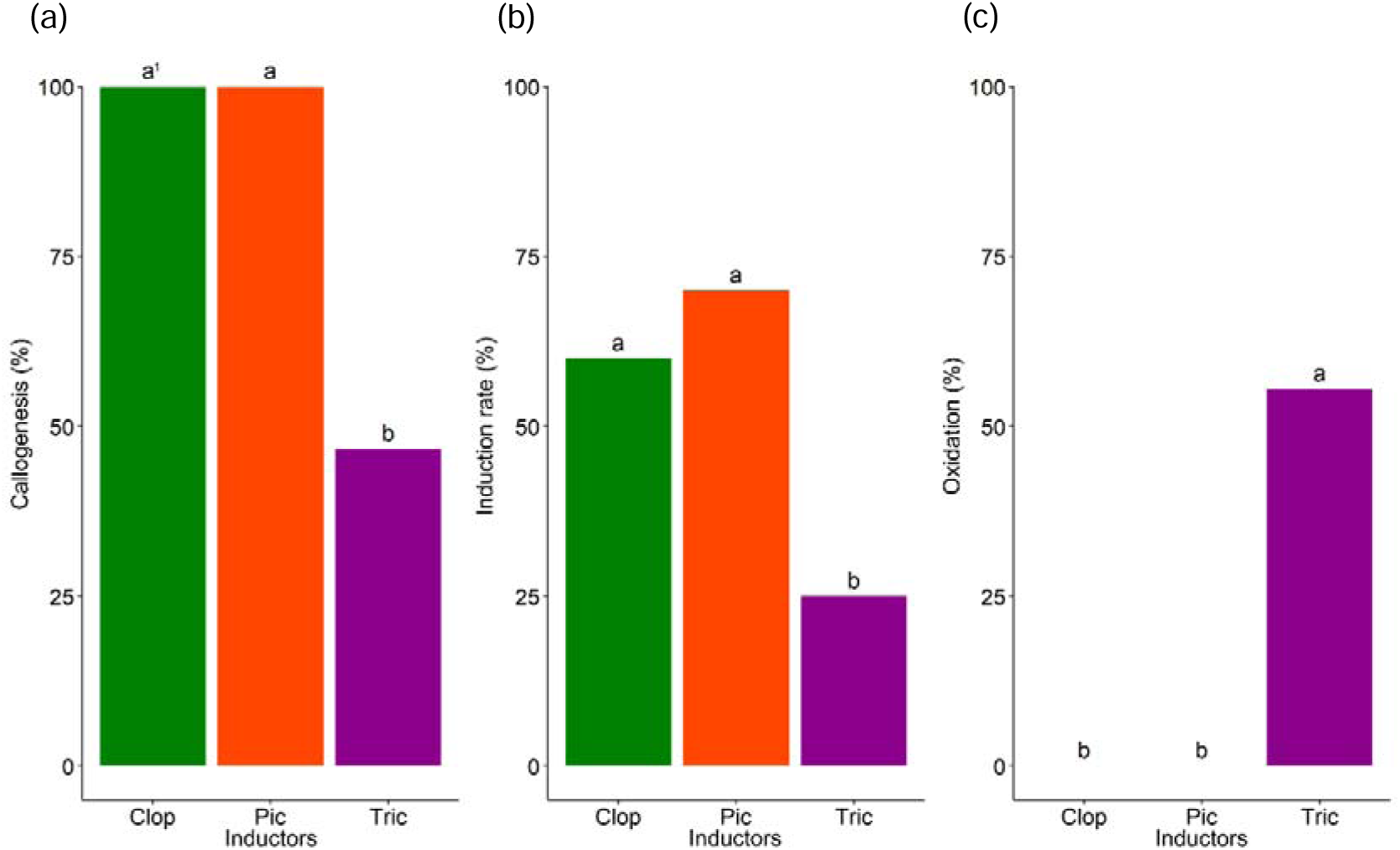
Non-embryogenic and embryogenic calli obtained upon induction of *Euterpe edulis* with picloram (150 µM). (a–h) Non-embryogenic opaque white rough callus (a, b), white spongy rough callus (c, d), smooth white callus (e, f), and smooth yellow callus (g, h). (i, j) Embryogenic rough yellow callus with somatic embryos (blue arrows). Scale bar: 2000 µM (a, c, e, g), 1000 µM (b, d), 500 µM (f, h, i), and 200 µM (j).

Using AFM, it was possible to differentiate the topography, profile, and elasticity of each callus type, thereby enabling the generation of a roughness pattern (Fig. 12). Opaque white rough calli displayed the lowest peak-to-peak (586.933 nm) value (Fig. 12a–d). White spongy and smooth white rough calli were similar to each other, with peak-to-peak values of 1269.52 and 1084.00 nm, respectively, predominance of valleys (SSK < 0), and similar symmetry (Fig. 12e–l). The smooth yellow callus was the most symmetrical (SKU ≈ 3), with greater balance between peaks and valleys (Fig. 12m–p). In spite of such differences, these patterns differed markedly from that of an embryogenic callus (yellow rough), which was characterized by the highest peak-to-peak value (4252.07 nm), a predominance of peaks, and the strongest asymmetry (SKU = 7.2885) (Fig. 12q–t).

**Fig. 12.**
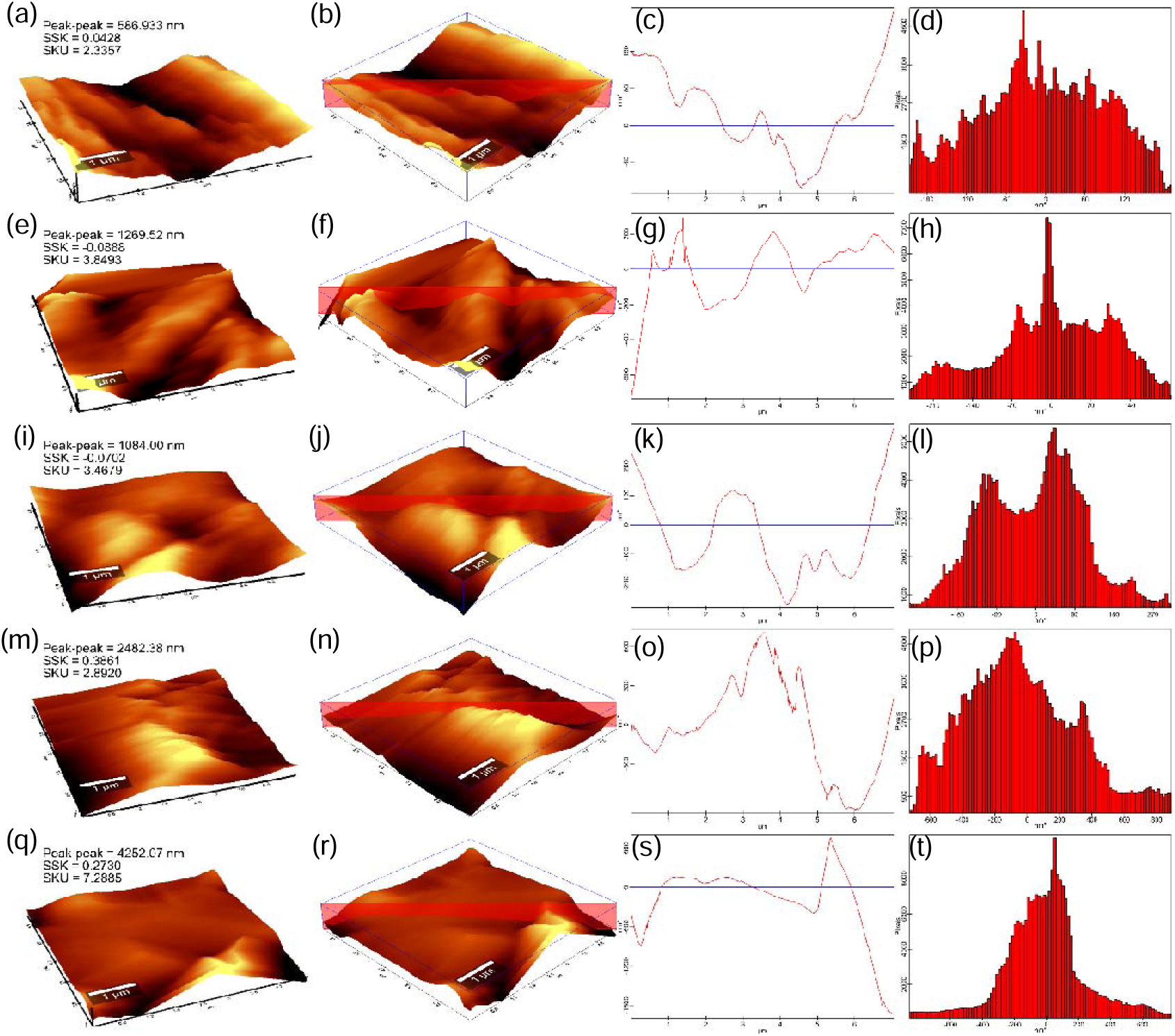
AFM images of the callus surface based on different calli types and analyses. (a–p) Non-embryogenic calli: rough white opaque callus (a–d), rough white spongy callus (e–h), smooth white callus (i–l), and smooth yellow callus (m–p). (q–t) Embryogenic rough yellow callus. The following analyses are presented: (a, e, I, m, q) topography, (b, f, j, n, r) cross-sections, (c, g, k, o, s) graphs of cross-sections, and (d, h, l, p, t) histograms of the topography.

Based on phase data obtained by AFM, the variation in the elasticity of each callus was determined (Fig. 13). Among non-embryogenic calli, the greatest phase variation occurred in the opaque white rough callus (Fig. 13a, b) and the least in the embryogenic callus (Fig. 13i, j), with the remaining specimens falling in between these two extremes (Fig. 13c–h).

**Fig. 13.**
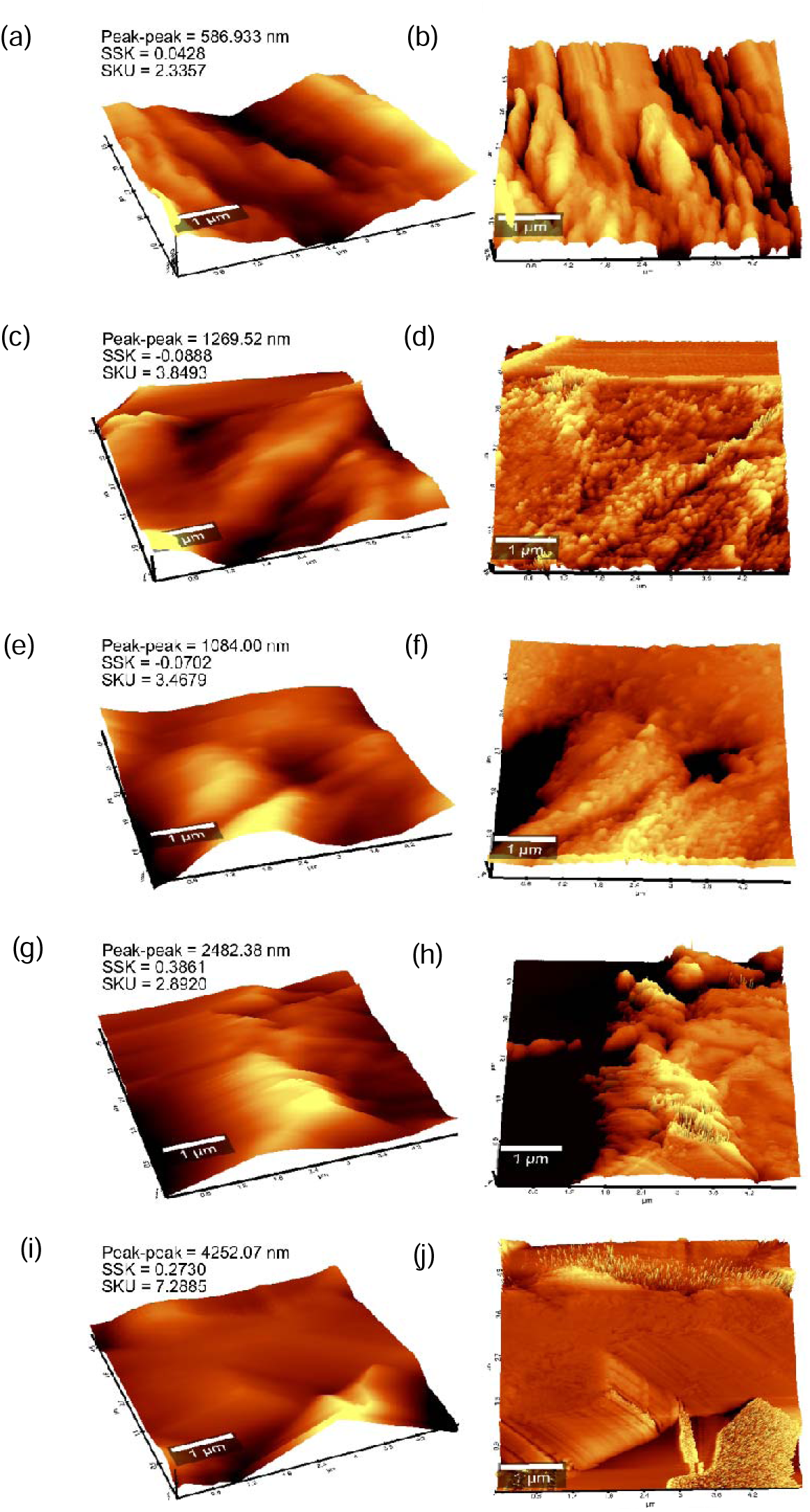
AFM images of callus surface topography. (a–h) Non-embryogenic calli: opaque white rough (a, b), spongy white rough (c, d), white smooth (e, f), and yellow smooth (g, h). (i, j) Embryogenic rough yellow callus.

### Chemical analysis of inducers

Based on PCA, zygotes induced with picloram and triclopyr clustered together (Fig. 14a). These results indicated that the two inducers shared a similar molecular structure and biological activity during embryogenic induction.

**Fig. 14.**
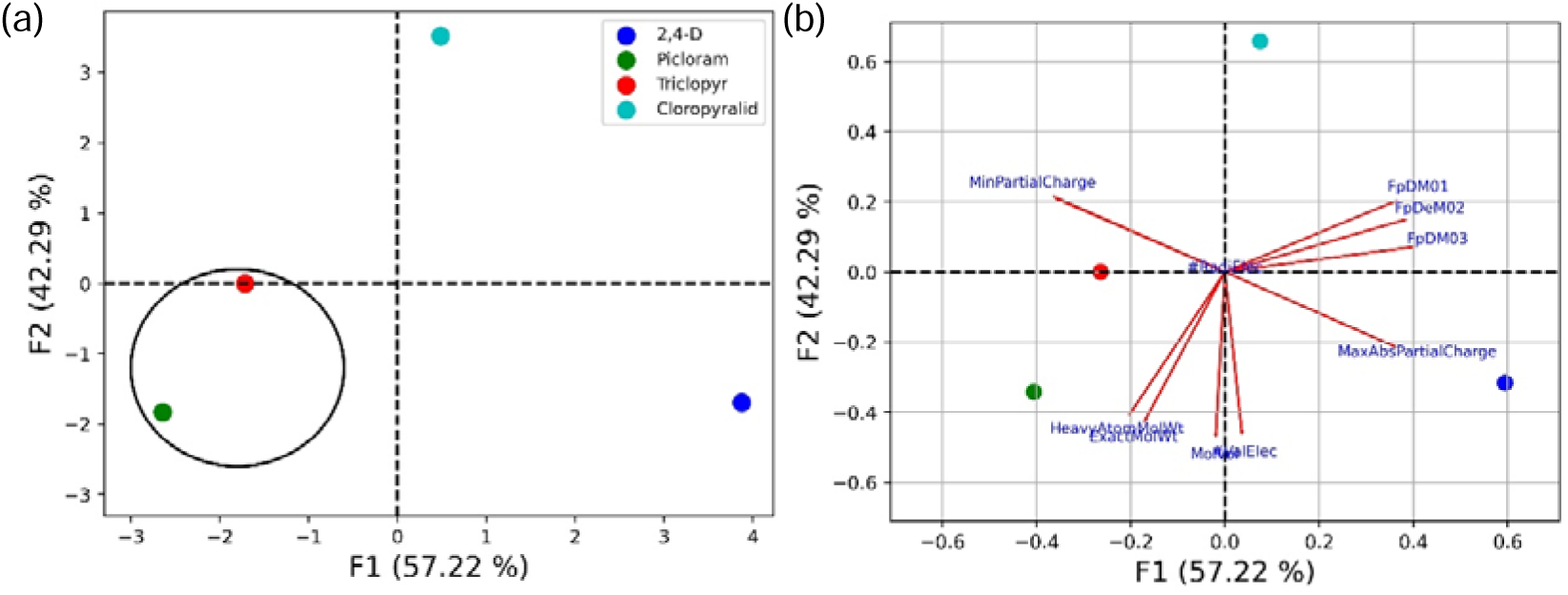
PCA of the embryogenic inducers 2,4-D, picloram, triclopyr, and cloropyralid. (a) Correlation graph of the first (F1; 57.22%) and second (F2; 42.29%) components of embryogenic inducers. (b) Scatter plot of the molecular properties obtained from PCA as a function of components 1 (F1) and 2 (F2).

Projections of the molecular properties corresponding to the first two principal components, F1 (57.22%) and F2 (42.49%), revealed a positive correlation between MaxAbsPartialCharge, MaxPartialCharge, and MinPartialCharge, but a negative correlation between FpDM01, FpDM02, and FpDM03 (Fig. 14b). A positive correlation was observed also for HeavyAtomMolWt and ExacMolWt, as well as ValElect and MolecVol. MaxAbsPartialCharge and MaxPartialCharge descriptor features were used to create the PCA model, which is summarized as a heat map for the first PCA components and the descriptor or characteristics used to statistically represent 2,4-D picloram, triclopyr, and clopyralid (Fig. 15).

**Fig. 15.**
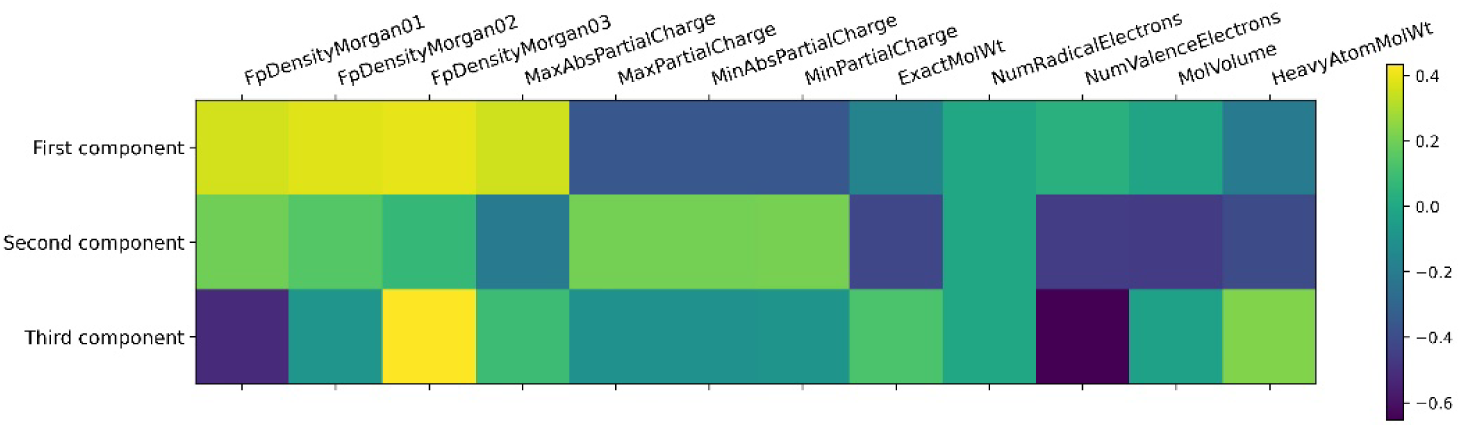
Heatmap representiing the correlation graph of the three components from PCA, in which F1 (57.22%) and F2 (42.29%) were the two main components, with the various descriptors.

### Maturation and germination

No visible changes were detected in picloram-induced embryogenic calli during maturation. However, a more detailed analysis revealed cell plasmolysis and features pointing to cell death, such as few organelles, rare mitochondria, lysosomes, amyloplasts, endoplasmic reticulum, and cytoplasmic degeneration (Fig. 16).

**Fig. 16.**
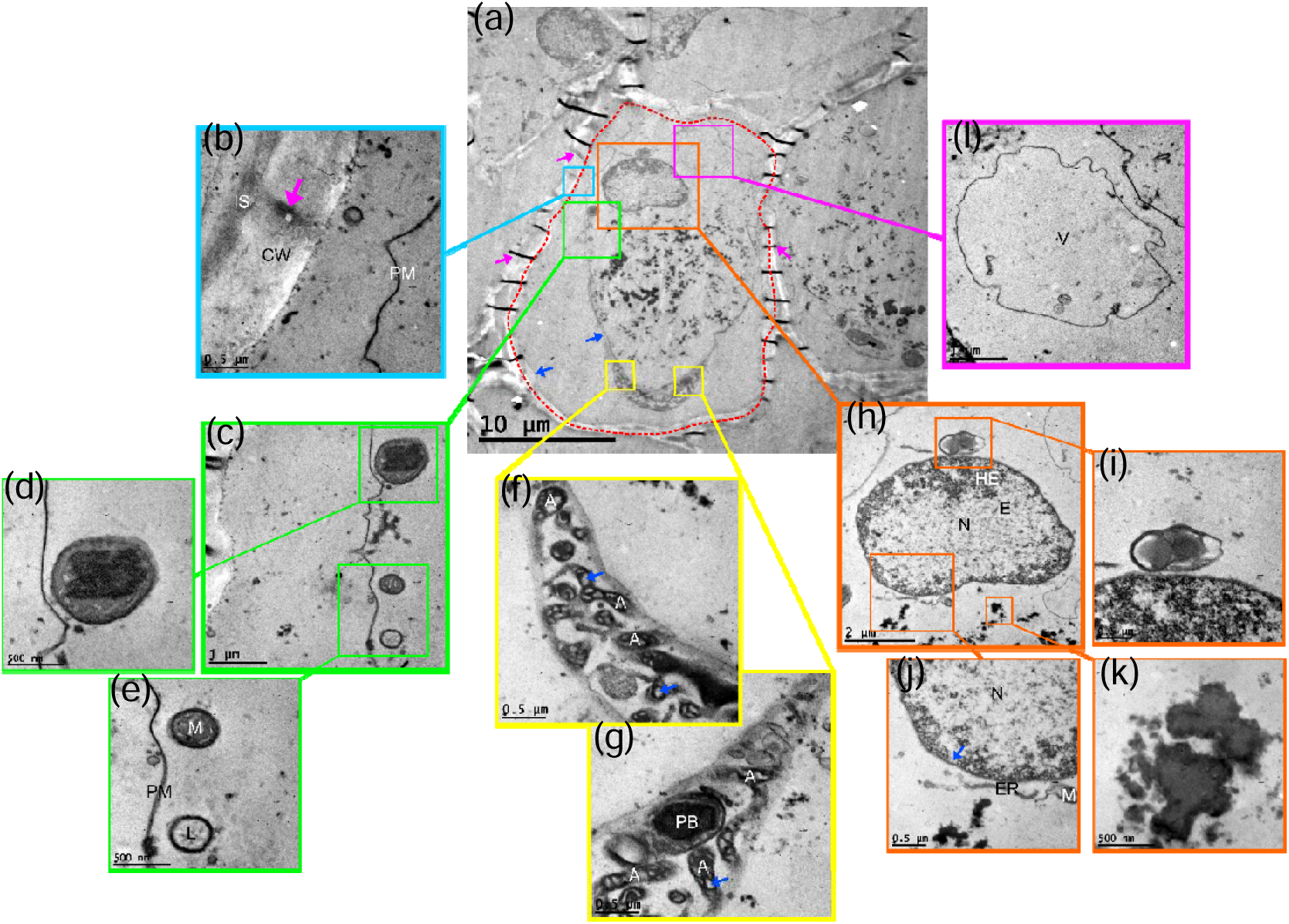
Ultrastructural characterization of embryogenic calli of *Euterpe edulis* induced with picloram (150 µM). (a) Cell (dashed red line) of embryogenic calli at maturation, with visible cell wall, plasma membrane (blue arrows), and plasmodesmata (pink arrows). (b–l) Details of the cell represented in panel (a): cell wall (CW), intercellular space (IS), plasma membrane (PM), and plasmodesma (pink arrow) (b); organelles close to the plasma membrane (c); plastid (d); mitochondria (M) and lysosome (L) (e); amyloplasts (A) and starch granules (blue arrows) (f, g); protein body (PB) (g); nucleus (N), heterochromatin (HE), euchromatin (E) (h); plastid (i); endoplasmic reticulum (ER), nuclear membrane (blue arrow) (j); cytoplasm degeneration (k); and vacuole (V) (l).

Calli obtained following induction with triclopyr (100 µM) displayed a strong increase in somatic embryos (+153.44%) when transferred from induction medium to pre-maturation (basal medium) for 30 days (Fig. 17a, b and Fig. 18a). However, the somatic embryos remained asynchronous and we were able to observe embryos at different stages (globular, scutellar, and coleoptilar) and sizes (Fig. 17c).

**Fig. 17.**
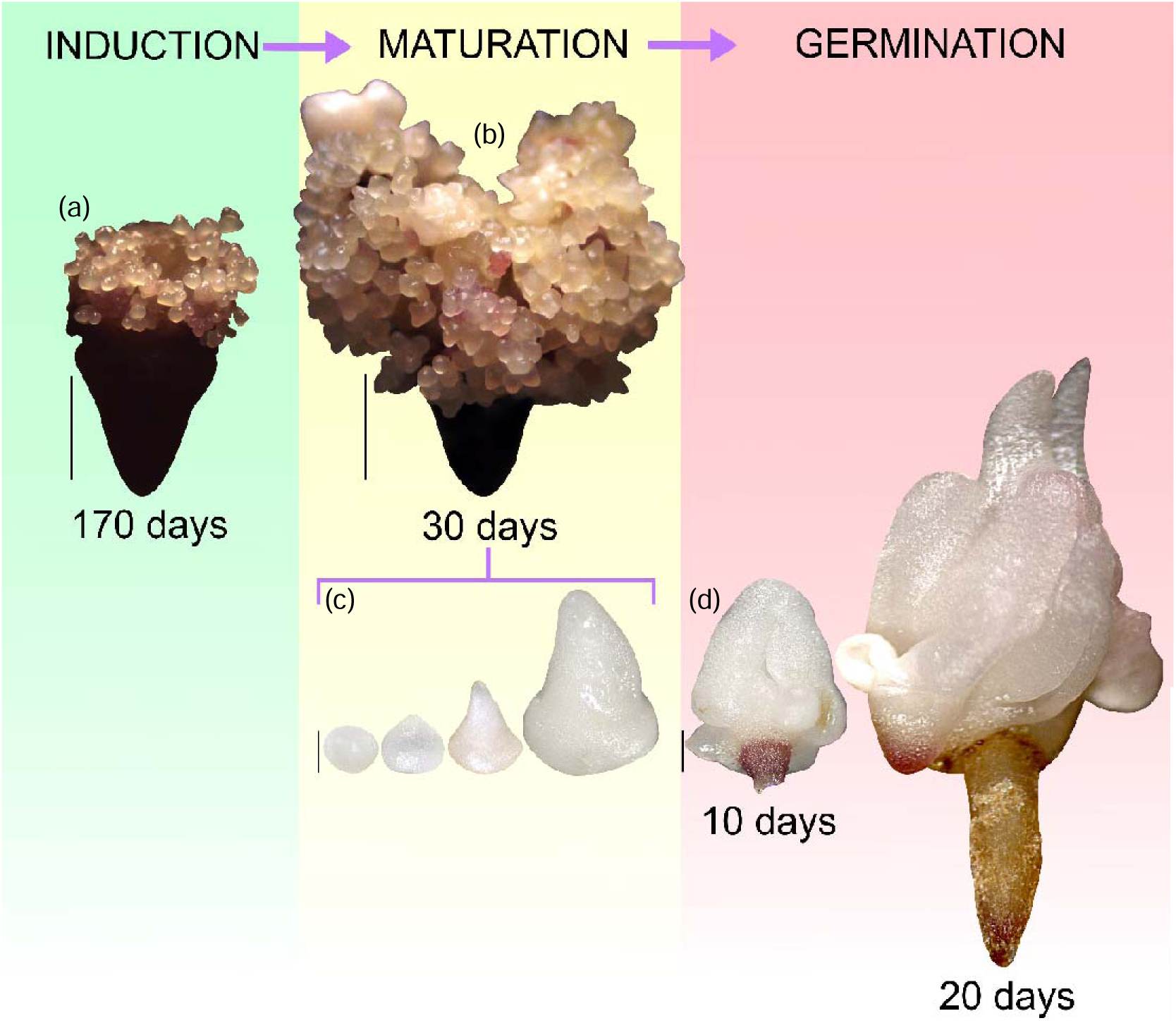
Embryogenic callus induced with triclopyr (100 µM) at different stages. (a) Induction stage with emergence of somatic embryos. (b, c) Embryogenic callus at maturation after 30 days in basal culture medium (b) with asynchronous somatic embryos (c). (d) Somatic embryo at germination after 10 and 20 days. Bar: 2000 µm (a, b) and 500 µm (c, d).

**Fig. 18.**
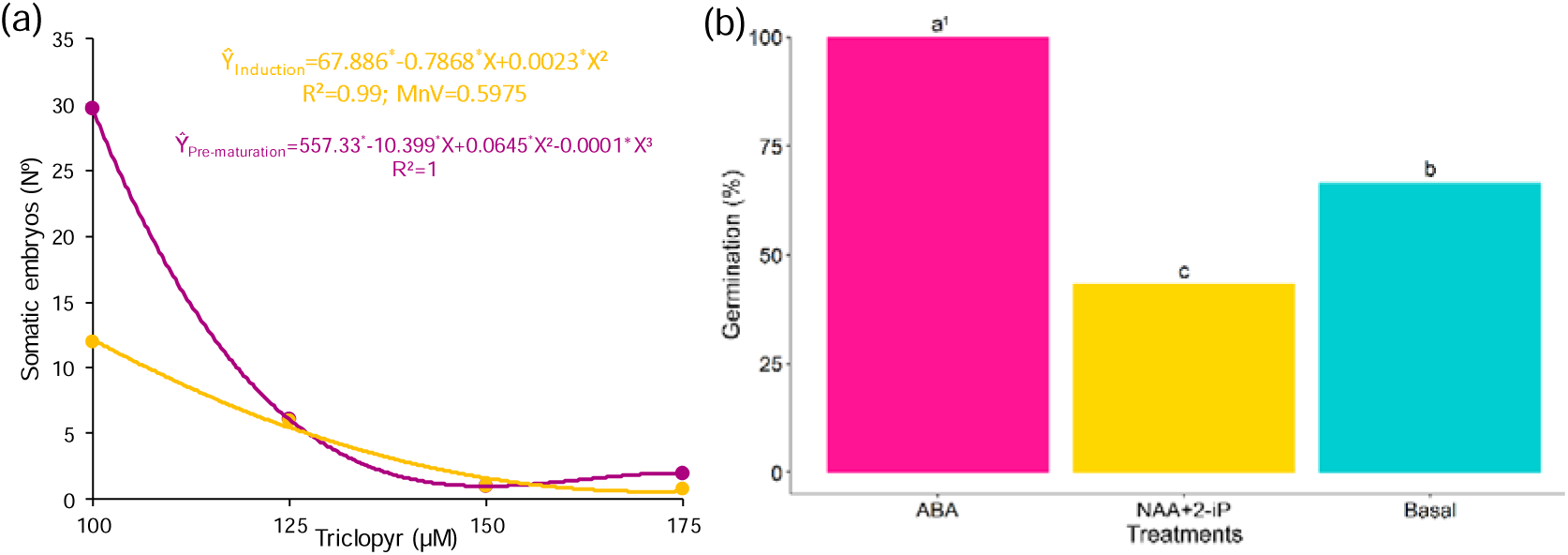
Number and germination rate of somatic embryos of *Euterpe edulis* treated with triclopyr, ABA, and NAA+2-iP. (a) Number of somatic embryos following induction with triclopyr at different concentrations (100, 125, 150, and 175 µM) measured at induction (yellow line) and after 30 days in basal culture medium (purple line). (b) Germination of somatic embryos induced with triclopyr (100 µM) and cultivated under different maturation conditions (ABA, NAA+2-iP, and basal culture medium). *Statistically significant coefficient at 5% by the *t*-test. ¹Means not followed by the same letter differ by Tukey’s test (*P < 0.05*).

Somatic embryos began to germinate after 10 days in culture medium supplemented with ABA and NAA+2-iP, as well as in basal medium. After 20 days, it was possible to observe the formation of normal somatic embryos, characterized by well-developed aerial parts and root systems (Fig. 17d). Somatic embryos obtained upon cultivation in medium supplemented with ABA achieved the highest mean germination rate (100%) (Fig. 18b).

### Growth and acclimatization of somatic emblings

Following the transfer of newly germinated somatic emblings to basal medium in test tubes, the highest survival was observed in somatic seedlings originating from culture medium supplemented with ABA and NAA+2-iP (67.58% and 60.32%, respectively), and the lowest in those grown in basal medium (69.84%) (Fig. 19a).

**Fig. 19.**
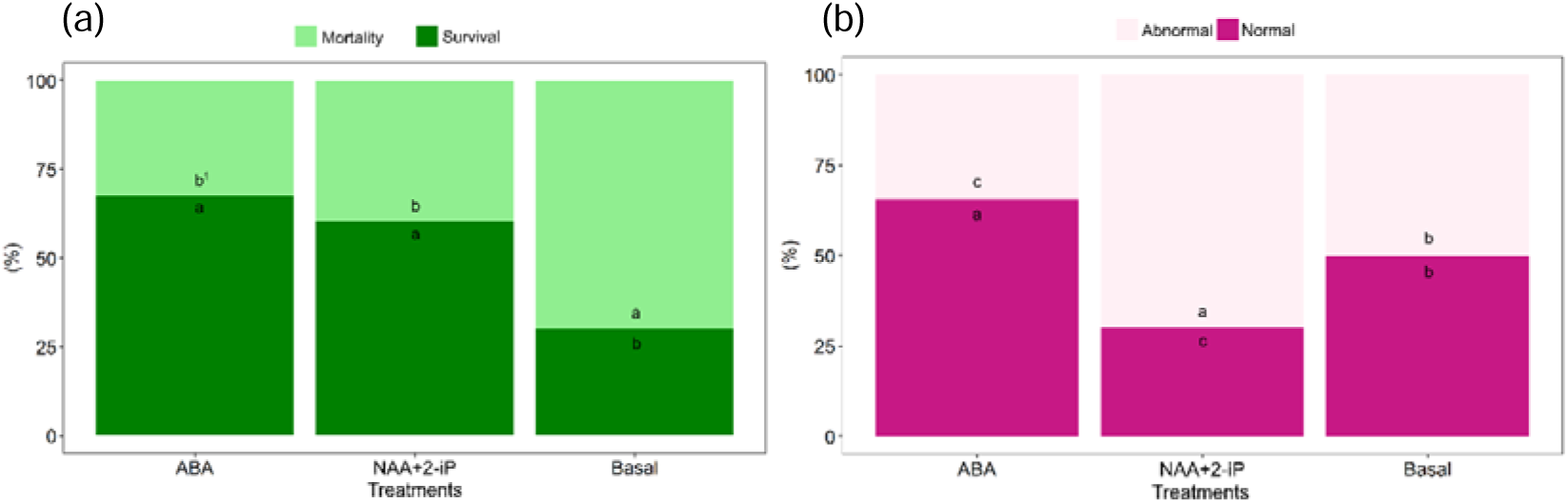
Charactteristics of somatic emblings obtained upon induction with triclopyr (100 µM) and subjected to different maturation conditions (ABA, NAA+2-iP, and basal medium). (a) Mortality and survival of germinated somatic embryos. (b) Percentage of abnormal and normal somatic emblings. ¹Means not followed by the same letter differ by Tukey’s test (*P < 0.05*).

Somatic emblings emerging from culture medium supplemented with ABA presented the highest percentage of normal somatic seedlings (65.48%) (Fig. 19b). In contrast, those maintained in culture medium supplemented with NAA+2-iP were characterized by the highest percentage of abnormal seedlings (70%) (Fig. 19b), with somatic seedlings presenting a thick collar and/or no shoot or root formation. Somatic emblings remained green throughout the acclimatization process (Fig. 20).

**Fig. 20.**
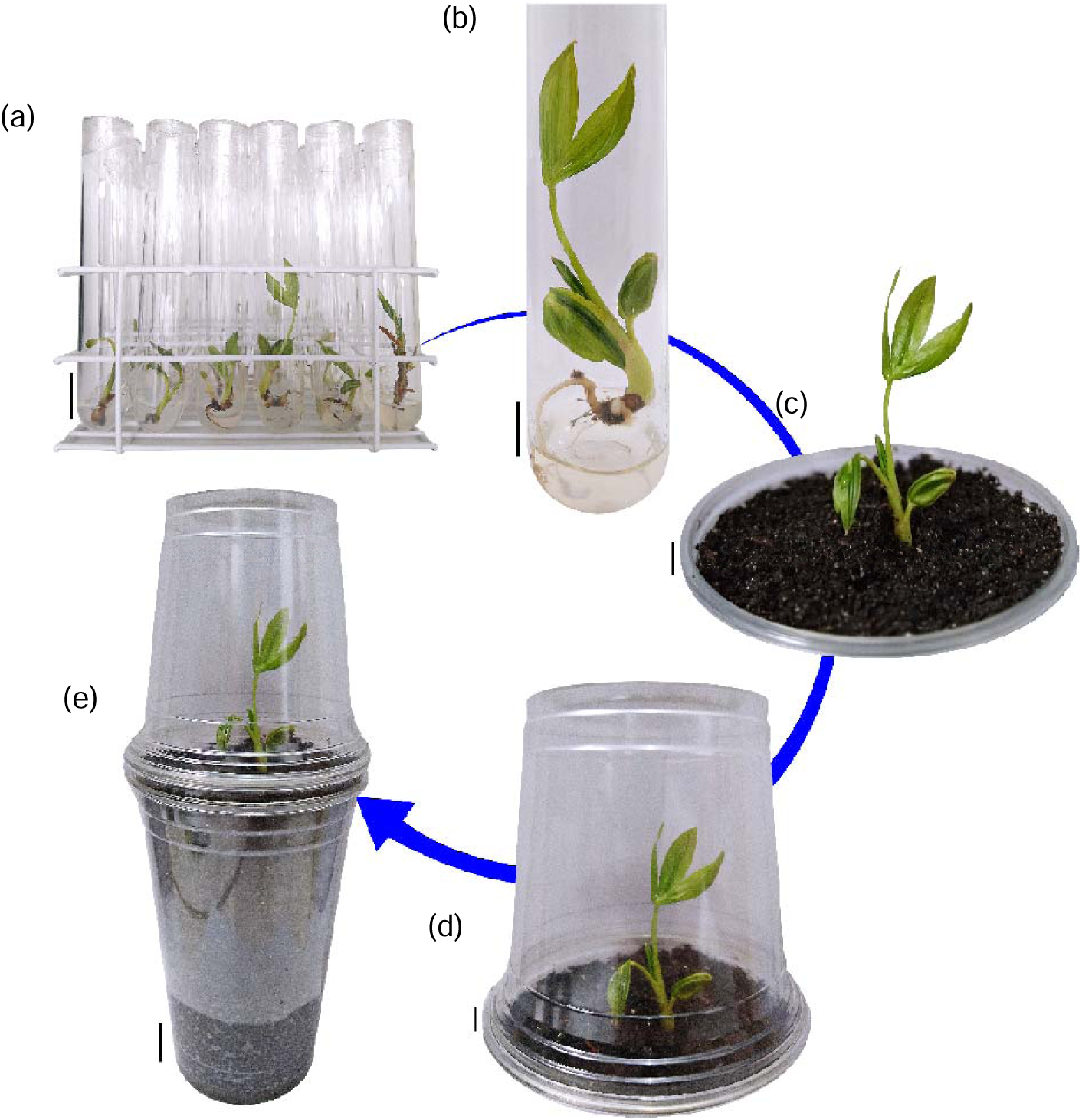
Acclimatization of emblings obtained from embryos induced with triclopyr (100 µM) and subjected to different maturation treatments (ABA, NAA+2-iP, and basal medium). Bar: 1 cm (b–d) and 2 cm (a, e).

## DISCUSSION

Since the pioneering work on somatic embryogenesis of *E. edulis* by Guerra and Handro (1988), further studies have followed (Guerra and Handro, 1988; 1998; Ledo *et al*., 2002; Mello *et al*., 2023; Oliveira *et al*., 2022; Saldanha *et al*., 2006; Saldanha and Martins-Corder, 2016). Here, we report the low efficiency of 2,4-D but high efficiency of picloram (150 µM) as auxin inducers of somatic embryogenesis in *E. edulis*. The scarce effectiveness of 2,4-D can be explained by its tendency to oxidize plant tissues, as observed also by Ferreira *et al*. (2022a) in the zygotic embryos of *Euterpe precatoria*, where picloram was 3.7 times more efficient than 2,4-D.

In the M9-10a strain of *Medicago truncatula* Gaertn., 2,4-D is present in low amounts during the induction phase, but is required for embryogenic callus formation and embryonic development (Orłowska and Kępczyńska, 2020). At higher concentrations, 2,4-D promotes the accumulation of O_2_, which interferes with the formation of calli and embryos. It also promotes the expression of the genes CLF, MSI1, FIE, and VRN2 involved in forming the PRC2 complex and, consequently, callus and embryonic development in *M. truncatula*. Oliveira *et al*. (2022) attained superior results with picloram (300 μM, 76.33 embryos) compared to 2,4-D (159.42 μM, 44.33 embryos) during somatic embryogenesis of *E. edulis*.

The presence of meristematic centers is a hallmark of embryogenic calli, and the formation of proembryos can be detected based on the development of these centers. Their formation is related to the response to hormonal signals (Freitas *et al*., 2016). Meristematic centers undergo active division, leading to the formation of small, compact cells with dense cytoplasm and thick walls (Oliveira *et al*., 2022), as was observed here in histological and transmission electron micrographs.

The first tissue that can be identified during embryogenesis is the protoderm, the outermost uniseriate layer formed by cells with a predominantly rectangular shape and centralized nuclei (Ferreira *et al*., 2022a, b). Subsequently, procambial cords composed of elongated cells with evident nuclei are formed (Ferreira *et al*., 2022a).

Another important feature is the multicellular origin of somatic embryos whereby they seem to merge with the tissue of origin, along with the fusion of somatic embryos (Freitas *et al*., 2016). The presence of idioblasts with inorganic inclusions (raphides) of acicular calcium oxalate crystals is common in palm trees and has been reported in the embryos of *E. oleracea* (Menezes Neto *et al*., 2010), *E. precatoria* (Ferreira *et al*., 2020), and *E. edulis* (Oliveira *et al*., 2022).

Here, we used AFM to investigate embryogenic induction in calli. AFM does not require costly sample preparations, and provides detailed information on topography, roughness, and elasticity of a surface. Moreover, calli can be analyzed by AFM without any need for prior dehydration or drying steps. Using AFM-derived information on topography, profile, elasticity, roughness, and symmetry, it was possible to differentiate embryogenic from non-embryogenic calli. Specifically, embryogenic calli displayed a much more pronounced roughness and symmetry compared to non-embryogenic calli.

Chemically, 2,4-D is derived from phenol and is classified as a phenoxyalkanoate within the class of synthetic auxins. Instead, picloram, triclopyr, and clopyralid belong to the class of pyridinecarboxylates, which share a a common pyridinium ring and carboxyl group (Duke and Dayan, 2011). The presence of this carboxyl group may explain the effectiveness of these three inducers in somatic embryogenesis, while the nitrogen present in the ring may underscore the mechanism of action of picloram and its analogs.

Picloram and triclopyr possess three chlorine atoms; whereas 2,4-D and clopyralid contain only two. Triclopyr performed better than the other inducers during somatic embryogenesis of *E. edulis*. In *Carica papaya* L., 4-chlorophenoxyacetic acid achieved much higher embryogenic responsiveness than 2,4-D, indicating that the number of chlorine atoms was crucial for biological activity (Cipriano *et al*., 2018).

Triclopyr, a common herbicide, is quickly translocated in the plant, mainly through the symplastic route, which allows the selective absorption of solutes. It then accumulates in meristematic tissues (Cessna *et al*., 2002), which are poorly differentiated and/or undifferentiated, and contain intensely dividing cells with large nuclei and dense cytoplasm (Gomes *et al*., 2017; Meira *et al*., 2020), as noted here in cells treated with triclopyr (100 µM).

These events are in line with the concept of totipotent stem cells proposed by Verdeil *et al*. (2007), and with the observations by Silva-Cardoso *et al*. (2020) in somatic embryos. A high nucleus/cytoplasm ratio, large nuclei, prominent nucleoli, small vacuoles, and an abundance of organelles, (e.g., mitochondria, ribosomes, and endoplasmic reticulum) indicate intense RNA synthesis and high metabolic activity; hence, they are associated with ongoing embryogenesis (Silva-Cardoso *et al*., 2020; Stein *et al*., 2010).

While clopyralid induces auxin-like responses, its effect on RNA and protein synthesis is akin to that of 2,4-D (Turnbull and Stephenson, 1985). Therefore, among the three synthetic auxins tested, triclopyr emerges as a promising embryogenic inducer, especially for picloram-responsive species.

Pre-maturation medium (without regulators) was essential for the growth and development of somatic embryos, with a milky appearance at all stages of maturation within 30 days. It also favored the germination of embryos at more advanced stages, during the maturation process (after 10 days), and that of seedlings (after 20 days). Previously, *E. oleracea* cultivated for 30 days in differentiation and maturation media yielded only white and compact globular embryos, plus translucent somatic embryos whose appearance became milky only after 80 days (Freitas *et al*., 2016). In *E. precatoria*, incubation for 60 days in differentiation and maturation media, resulted in mostly elongated somatic embryos, with a whitish opaque color similar to that observed in the torpedo stage of development (Ferreira *et al*., 2022a).

During maturation and germination of somatic embryos in the present study, 100% of the isolated embryos achieved germination if grown in medium supplemented with ABA (5 µM), 67.48% of them survived, and 65.48% presented a normal appearance. This outcome was superior to that achieved when using NAA+2-iP or medium without regulators. The development of *E. edulis* embryos into somatic seedlings was first attempted using medium supplemented with NAA (2.68 µM) and 2-iP (24.6 µM), then adjusted to 0.53 µM NAA and 12.3 µM 2-iP (Guerra and Handro, 1988; 1989). The same approach was applied later on by Oliveira *et al*. (2022) and Scherwinski-Pereira *et al*. (2012), as well as Freitas *et al*. (2016) in *E. oleracea* and Ferreira *et al*. (2022a) in *E. precatoria*.

In most studies of somatic embryogenesis in *Euterpe* spp., the germination phase has not been quantitatively analyzed, and only the formation of normal somatic seedlings has been mentioned (Ferreira *et al*., 2022a; Guerra and Handro, 1988; 1998; Saldanha *et al*., 2006; 2012). The sole quantitative information came from Ledo *et al*. (2002) and Scherwinski-Pereira *et al*. (2012), who reported attaining 14.2 and 20.2 somatic embryos per callus, respectively; whereas Freitas *et al*. (2016) noted that 58.7% of somatic embryos were converted into seedlings.

Abnormalities in somatic embryos are common and are caused by genetic or epigenetic changes to DNA. These DNA changes can be influenced by external factors, such as the use of plant growth regulators and mutagenic substances, or stress factors, such as high and low temperatures, drought, salinity, and heavy metals (Garcia *et al*., 2019).

## CONCLUSION

The use of 2,4-D for somatic embryogenesis of *E. edulis* is not recommended as it is less efficient than picloram (150 µM) or its analog triclopyr (100 µM). The selection of somatic embryos can be improved through the use of AFM, which distinguishes embryogenic from non-embryogenic calli. Somatic embryos of *E. edulis* induced with triclopyr multiplied five times more rapidly if incubated for 30 days in pre-maturation (basal medium). The maturation and germination of somatic embryos of *E. edulis* was maximum in medium supplemented with ABA (5 µM). Emblings obtained through embryogenic induction with triclopyr displayed normal formation and acclimatization. The common chemical structure of picloram and triclopyr explains their similar biological activity as embryogenic inducers.

### CrediT authorship contribution statement

**Tamyris de Mello**: Investigation, Methodology, Writing, review and editing. **Yanara dos Santos Taliuli**, **Tatiane Dulcineia Silva**, **Tadeu Ériton Caliman Zanardo**, and **Clovis Eduardo Nunes Hegedus**: Investigation, Methodology. **Breno Benvindo dos Anjos**, **Edilson Romais Schmildt**, and **Adésio Ferreira**: Formal analysis. **Maicon Pierre Lourenço**, **Patricia Fontes Pinheiro**, **Glória Maria de Farias Viégas Aquije**, **José Carlos Lopes**, and **Wagner Campos Otoni**: Supervision, Writing – original draft, Writing – review and editing. **Rodrigo Sobreira Alexandre**: Conceptualization, Supervision, Project administration, Funding acquisition, Writing – original draft, Writing – review and editing.

### Funding

This study was funded by the Fundação de Amparo à Pesquisa e Inovação do Espírito Santo (FAPES) (EDITAL FAPES/CNPq N° 05/2017 - PRONEM - PROGRAMA DE APOIO A NÚCLEOS EMERGENTES, registered with SICONV under n° 794009/2013 and Process FAPES n° 72660945 with EDITAL FAPES N° 08/2021 – MANUTENÇÃO DE EQUIPAMENTOS) and Conselho Nacional de Desenvolvimento Científico e Tecnológico (CNPq) (Finance code 308365/2019-4).

### Data availability Statement

Data sharing is not applicable to this article as all new created data is already contained within this article.

## Acknowledgments

The authors would like to thank Conselho Nacional de Desenvolvimento Científico e Tecnológico (CNPq) and Fundação de Amparo à Pesquisa e Inovação do Espírito Santo (FAPES) for research funding, and Editage (www.editage.com) for English language editing. Acknowledgments are extended also to the Carlos Alberto Redins Cellular Ultrastructure Multiuser Laboratory (LUCCAR) on behalf of Prof. Dr. Breno Valentim Nogueira.

